# Reducing Glioblastoma Cell Aggressiveness via Static and Dynamic Magneto-Mechanical Stimulation with Vortex Microdiscs on Substrates of Physiological Stiffness

**DOI:** 10.1101/2025.10.15.682498

**Authors:** Andrea Visonà, Sébastien Soulan, Bernard Dieny, Robert Morel, Alice Nicolas

## Abstract

External mechanical stresses acting on cellular compartments critically regulate cell behaviour and can induce cell death. Magnetically actuated particles present a promising strategy to apply such forces in a controlled manner, with potential applications in cancer therapy. In this study, we investigate the effects of actuating vortex magnetic microdiscs on a glioblastoma cell line cultured on soft, biomimetic substrates that mimic in vivo stiffness. Using a Halbach array, we applied either static mechanical compression or a combination of compressive and low-frequency vibrational stresses (2–20 Hz). Our results demonstrate that both compressive and vibrational stresses impair important cellular functions associated with glioblastoma persistence in a dose-dependent and stiffness-dependent manner. In particular on soft 2D substrates, sufficiently strong compressive loads limit proliferation, while the addition of vibrations alter cell motility, cell morphology and the acto-myosin machinery. Our findings demonstrate that magnetic particles-mediated mechanical stimulation can disrupt glioblastoma cell aggressiveness in physiologically relevant 2D substrates, supporting its potential as an adjunct to conventional chemo-and radiotherapies by both inducing cell death and limiting resistant populations.

## 1 Introduction

Application of mechanical stresses on cellular compartments are known to influence cellular functions, being even able to cause cell death [1–3]. In this context, applying intracellular mechanical stresses is being considered as a potential adjunctive treatment in some pathologies. For instance, nuclear compression triggers change in cell migration mode [4], and the deformation of the membranes of the Golgi apparatus or of the endoplasmic reticulum affects the flow of calcium or other ions, with consequence on cell metabolism, polarity, migration, proliferation [1]. These forces may arise from the stimulation of intern agents, such as actin fibre contraction or osmotic imbalance [5]. They may also be caused by external stresses such as hydrodynamic flows. Recently, advanced microsystems based on optical or magnetic traps have been developed to remotely apply controlled mechanical stresses to precise locations in cells [6–8]. For example, magnetic particles have been used to apply traction forces on neurites and improve their outgrowth in the context of neural regeneration [9, 10]. Similar particles were set in vibration by an alternating magnetic field to stimulate insulin secretion in pancreatic cells [11] or calcium influx in mechanosensory neurons [12]. Several studies have also shown that magnetic particles set in vibration in the Hz range could induce cancer cell death [13–18], possibly by mechanically altering the plasma or the lysosomes membranes.

More specifically, the use of magnetically-actuated particles has opened perspectives for the cure of glioblastoma, a lethal cancer for which no curative treatment is currently available [19]. The challenge in curing glioblastoma resides in four main reasons: its invasiveness (aggressive migration behaviour), heterogeneity (both patient-to-patient wise but also intra-tumour wise), its rapid development and its resistance to chemotherapy [20]. In addition there is the difficulty of circumnavigating the brain blood barrier that, while protecting the brain and making it a highly controlled and exclusive environment, represents a real challenge for molecules to be effectively delivered to brain tumours. In a pioneering study, Kim et al. [13] showed that low frequency vibrations of magnetic microdiscs attached to the cell membrane could trigger apoptotic pathways in relation to the stretching of the plasma membrane and the activation of specific ion channels. Since then, several groups have shown that internalized particles put in vibration at a frequency of few tens of Hz could efficiently kill model glioblastoma cells [15–17]. In contrast to radiotherapy, the first-line treatment for glioblastoma, particle-based treatment may target specific cell populations by biofunctionalising the surface of the particles. However, it has been shown that killing cancer cells was not successful to eradicate glioblastoma, as the surviving cells drift toward more aggressive phenotypes [21]. For this reason, combination of treatments that both destroy cancer cells and limit the aggressiveness of the surviving ones are currently underway [22]. Regarding magnetic particles, once the mechanical forces they transmit have been optimised, they could be used as adjunctive treatment, enhancing the efficacy of primary therapy and limiting the activity of cells resistant, for example, to radiotherapy [23].

In the past studies, the in vitro demonstrations of the efficacy of treating cells with vibrating particles were carried out in standard culture plates, with plastic surfaces (see [24, 25] for a review). But the effectiveness of the vibrating particles to kill cancer cells was not as successful with cells grown as spheroids [25] or in vivo [15, 26]. A basic limitation of standard plastic plates is that, while they are highly effective in enabling cell growth and imaging, these culture conditions put cells in a mechanical environment bearing little or no resemblance to that met in vivo, for instance being a million times stiffer [27, 28]. This is of primary importance, since the mechanical properties of the extracellular environment have been shown to be a key determinant of invasion, proliferation, and resistance to radio-chemotherapy in glioblastoma [29–31]. Although the difference between the in vitro and the in vivo assays cannot be solely attributable to the difference in mechanical properties of the extracellular environment, the interaction of the magnetic particles with cells, as well as the signalling pathways involved in cellular functions such as proliferation or migration, are regulated differently under these culture conditions [32–34]. Standard culture conditions are for instance known to induce drifts of cell phenotypes [35], while cells grown either on soft substrates mimicking the stiffness of the tissue of interest or in 3D as spheroids were shown to better preserve cellular phenotypes [36, 37].

In this context, we questioned whether mild stresses applied by internalized particles could impair some basic cellular functions involved in glioblastoma aggressiveness, for instance cell migration and proliferation, when the cells display a physiologically relevant level of contractility. To this end, a model glioblastoma cell line was grown in vitro on soft supports whose stiffness matches that of glioblastoma tissues to achieve a relevant mimicry of the cellular characteristics encountered in vivo. These simple substrates provide an easy platform to optimise the intracellular forces generated by the particles, before moving to more advanced in vitro systems. Following previous studies from our group and others, we used magnetically actuated vortex microdiscs (MDs) to apply the mechanical stresses [24]. Vortex microdiscs are highly anisotropic particles in which the magnetisation in the absence of field has a vortex configuration. This configuration limits the remanent magnetisation of the particles and enables them to remain dispersed. Alternating or rotating magnetic fields with a frequency of few Hz or tens of Hz were shown to set the MDs in vibration or rotation. This treatment was shown to trigger cell death in vitro in a dose-dependent manner. In addition, it was shown that increasing the frequency had a decreasing impact on cell viability, suggesting that vibrations in the Hz regime may have a stronger impact on cell fate compared to tens of Hz [13]. This raises the question whether a static magnetic stimulation of the MDs, resulting in a static mechanical force may also alter cell functioning. In this context, we analysed the impact of intracellular mechanical stimulations consisting of either a static mechanical stress or the superimposition of static and dynamic vibrational stresses in the range of 2 to 20 Hz, all mediated by the MDs.

In this article, we thus report on the the sensitivity of U87-MG cells to compressive and vibrating stresses resulting from the actuation of magnetic microdiscs cells, highlighting a dose-dependent effect and a sensitivity on the elastic properties of the extracellular environment, the more pronounced impact being observed in cells grown on soft, biomimetic substrates. We show that gentle mechanical stimulation, obtained with a minimal dose of particles, is impairing cell proliferation and motility, while irreversible alteration of the cells shape is observed with particles vibration between 10 and 20 Hz. Furthermore, we observe that sufficiently large static compressive forces can impair cell proliferation, although vibrating mechanical stresses have a more pronounced effect, with increased persistence at higher frequency. The impairment of cell proliferation and motility is attributed to alterations of cell vimentin and actin cytoskeletons, that are shown to drive changes in cell contractile capabilities. By showing that intracellular mechanical stimulation comprising compression and vibration can both induce cell death and limit cell contractility, proliferation and migration of resistant cells, our study therefore highlights the potential of particle-mediated mechanical therapy in being effective in the treatment of cancer as an adjunct to chemotherapy or radiotherapy.

## 2 Methods

### 2.1 Magnetic microdiscs fabrication and characterization

The vortex micro-discs were fabricated using a top-down approach as previously described [38]. Briefly, a double layer of PMMA positive resist and ma-N negative resist was deposited onto a 100 mm silicon wafer and patterned using deep ultraviolet optical lithography to create an array of circular wells with a diameter of 1.3 *µ*m in the ma-N resist. After developing the ma-N resist, three successive metal layers – gold (10 nm), permalloy (Ni_80_Fe_20_, 60 nm), and gold (10 nm) – were deposited using electron-beam evaporation (Fig. 1a). A first ethanol liftoff allowed removing the top ma-N resist, leaving the array of particles attached to the base PMMA resist (Fig. 1b). The magnetic properties of the particles were characterized using vibrating sample magnetometry (MicroSense EasyVSM). Figure 1c shows the magnetization curve with in-plane applied magnetic field, indicating a 60 mT saturation field and a very low magnetic remanence. As already reported, the low remanence ensures that the particles do not aggregate when they are released in a liquid [38]. On the other hand, due to their low saturation field, the particles can be remotely actuated using permanent magnets devices. A second acetone lift-off allowed to detach the particles from the PMMA base resist (Fig. 1d). Particles were sonicated three times with hot acetone to remove any resist monomer, after which they were washed three more times with ethanol and stored in ethanol. The geometric and structural specifications of the free vortex particles were obtained by SEM imaging as described in [32]. Prior to use, particles were dispersed by sonication and the ethanol was replaced with culture medium, with extensive rinsing steps to minimize cell exposure to ethanol.

**Figure 1:**
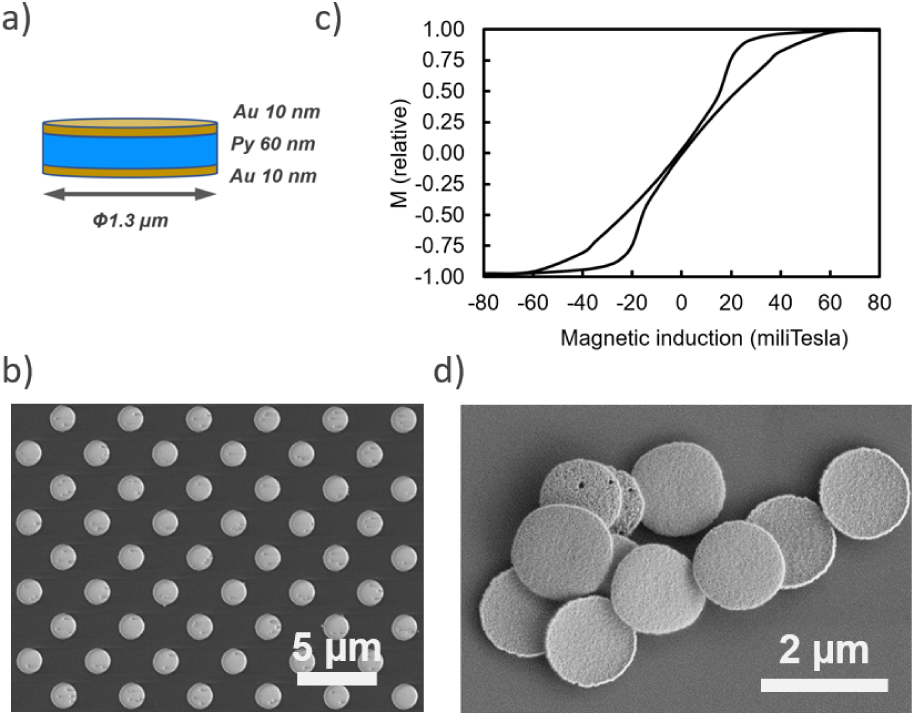
a) Schematics of the structure of vortex microdiscs. b) Microdiscs array still attached to the PMMA resist, after lift-off of the ma-N resist. c) Magnetization curve of the still attached microdiscs with an in plane magnetic field. d) Liberated microdiscs, after lift-off of the PMMA resist.

### 2.2 Magnetic field device

The magnetic field was generated by an array of seven rare-earth magnets (NdFeB N42 magnets Supermagnete, Gottmadingen, Germany) assembled in a planar Halbach configuration [39]. The array is attached to an orbital stirrer (IKA MS3 Digital, Staufen, Germany) and set into motion under a fixed cell culture plate (Fig. 2a–b). The size of the magnets is 40 x 8 x 4 mm^3^, with the magnetisation either parallel to the horizontal *x* axis or the vertical *z* axis according to the position in the Halbach array. The spacing between each magnet is of 3 mm. The width and spacing of the magnets is optimised such that, over the center region of the array: 1) the field amplitude is almost constant at a distance of 4 mm above the magnets and 2) given the orbital amplitude of the stirrer rotation, of 4.5 mm, the angular variation in the field direction at a fixed point above the array is maximum (Fig. 2c–d). As it is, the device allows for an oscillating magnetic field of 250 mT, with an angular amplitude of 55 degrees and a frequency up to 25 Hz. The device is operated inside an incubator at 37*^◦^*C with controlled CO_2_ and humidity levels.

**Figure 2:**
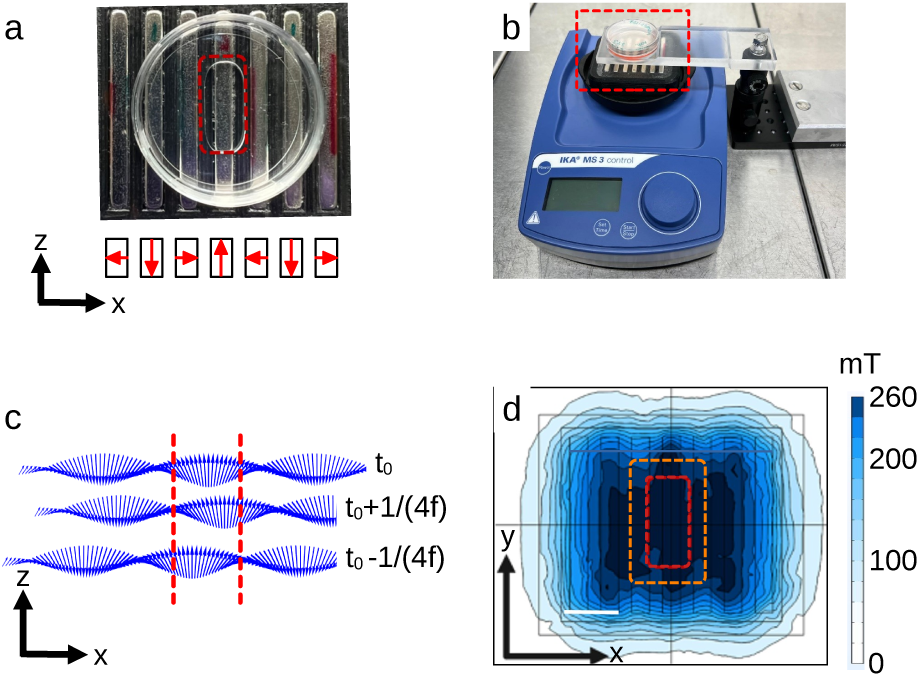
a) Top view picture of the rare-earth magnets array in a planar Halbach configuration, with a 35 mm diameter Petri dish. The red dashed rectangle indicates the zone of the dish used for cell culture. The schematics at the bottom shows the magnets magnetisation direction, with *z*-axis being out-of-plane. b) The orbital stirrer with the magnets array and the suspended Petri dish. The magnitude of the magnetic field is adjusted through the height of the beam. c) Side view of the field vector, 4 mm above the magnets, when the array is set in rotation. The orientation of the field is shown in the static condition (t_0_) and for the largest displacements of the magnets along the *x*-axis, at t_0_± 1/(4f), with f the oscillation frequency. d) Norm of the oscillating field (Bar: 1 cm). In (c) and (d), cells within the red dashed lines, standing still above the array, are exposed to an oscillating field having a nearly constant norm. The orange dashed rectangle shows the amplitude of the orbital oscillation.

### 2.3 FEM modelling of the field and magnetic forces and torques

Finite element method modelling of the magnetic field created by the planar Halbach array was performed using the COMSOL Multiphysics^®^ software (COMSOL AB, Stockholm, Sweden, v. 6.3). The force **F** on a MD with magnetisation **m**, resulting from the magnetic field **B** gradient, is given by

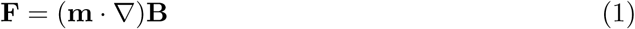

In this calculation, we made the assumption that the magnetisation is saturated in the direction of the local field:

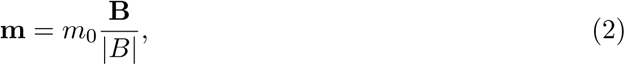

with *m*_0_ = 6.4e-14 *A* ∗ *m*^2^ [40]. The magnetic torque **T** acting on the particle is given by

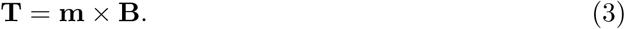

For this second calculation we have considered the three situations where **B** is alternatively applied along the *x*, *y* or *z* direction.

### 2.4 Design of culture substrates

Culture supports were purchased from Cell&Soft. The soft supports, of 10 kPa stiffness, consist of a polyacrylamide hydrogel layer covalently attached to a glass coverslip, and surface-coated with 1 *µ*g/cm^2^ of fibronectin. Glass supports were provided with an identical coating of fibronectin. 10 kPa substrates containing fluorescent beads of 200 nm were also purchased to performed traction force microscopy (Cell&Soft, Mecatract). The growth surface available on the substrate was limited to a specific area where the cells experience a constant field amplitude (Fig. 2).

### 2.5 Cell culture and magnetic stimulation

The cell line used in this study is a GFP-tagged U87-MG purchased from ATCC. Cells were grown in Dulbecco’s Modified Eagle’s medium (DMEM, Gibco, ref. 31966047) completed with 10% fetal bovin serum (FBS, Gibco, ref. 10270106) and 1% ATAM (Gibco, ref. 15240062). Cells were cultured in a humidified atmosphere of 5% CO_2_ at 37*^◦^*C. For the experiments, cells were seeded at a density of 4000 cell/cm^2^ on the soft and the stiff substrates. The microdiscs were introduced in the culture plates 6 hours post seeding at a concentration corresponding to 250 or 500 MDs/cell. The cells loaded with the magnetic particles were exposed to the magnetic field after 16 h of incubation. Cells were exposed to the field for 20 minutes, one condition right after the other. This consequently introduced a lag of 20 minutes between one and the other, which was taken into account during the cell assays. The field stimulation was performed at 37*^◦^*C in a humidified atmosphere of 5% CO_2_. After the magnetic stimulation, the culture medium was changed to remove any detached cells.

### 2.6 Quantification of the number of particles per cell

The number of particles interacting with cells was quantified at three different nominal concentrations on the polyacrylamide and glass substrates via vibrating sample magnetometry, following the protocol described in Ref. [32]. A linear correlation between the areal concentration of particles and the number of particles interacting with cells was found in the range of concentration studied (Fig. 3). The fitting function was used to adjust the nominal concentration to be added in the culture medium to obtain the desired amount of particles per cell. Two particle loads were chosen for the experiments, of 250 and 500 MDs/cell, which correspond to a areal concentration of respectively 5 and 10 *µ*g/cm^2^ for cells grown on the soft substrate, and of 10.3 and 15.5 *µ*g/cm^2^ for cells grown on glass.

**Figure 3:**
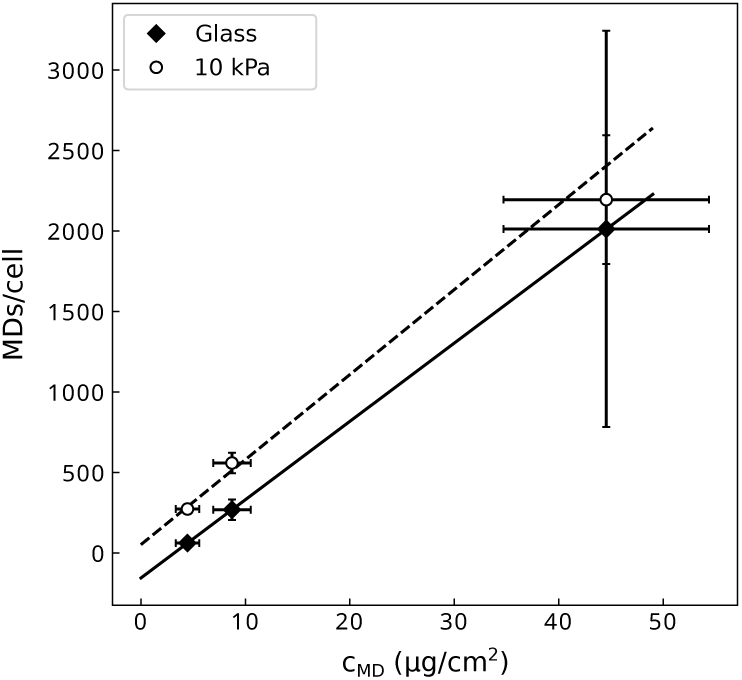
Relation between the concentration of microdiscs (MDs) delivered in the culture medium and the number of microdiscs interacting with the cells.

### 2.7 Cell growth, death and motility assays

Cell counting and tracking was performed by live staining the nuclei with Incucyte Nuclight Rapid Red Dye (Sartorius, ref. 4717). The DNA stain was used at a dilution of 1:2000. It was added concomitantly with the magnetic particles. Cell death was quantified using propidium iodide at a concentration of 1:2000 (PI, Molecular Probes, ref. P3566).

After the magneto-mechanical stimulation, the cells were transferred to an inverted microscope equipped with a motorised stage (IX83 Olympus, Olympus France, Rungis, France) and an incubation chamber with controlled temperature, humidity and CO_2_ pressure (Okolab, Rovereto, Italy). The cells were imaged at 10x magnification in five fields of view evenly distributed across the culture plate. Cell nuclei as well as PI-positive nuclei were counted 4 h and 24 h after the magneto-mechanical stimulation. Cell morphology was also quantified at these time points by capturing the green fluorescence of the cytoplasm. Cell motility was quantified by acquiring one image every 20 minutes for 3 h, beginning either 4 h and 24 h after the magneto-mechanical stimulation. The 20-minute time lag between samples imposed by the magnetic stimulation was taken into account in the implementation of the video time lapse. Since the time step is equivalent to the delay between one stimulation and the other, by carefully choosing the start and end frame of the three-hour sequence, we were able to extract a 2-hour time lapse that starts at the scheduled time of 4 h and 24 h for each condition. A control condition corresponding to cells with MDs not subjected to field was included for both type of substrates.

### 2.8 Image analysis

The nuclei of Nuclight positive cells were segmented with a trained Cellpose model [41]. The training was implemented starting from cyto2 pre-trained model as described in Ref. [32]. The cell number was obtained from the segmented masks with an in-house Python 3 script, summed over all fields of view, and normalized to the control condition, consisting of cells loaded with MDs but not exposed to the magnetic field. Given the low number of PI positive cells and the low contrasted signal, CY3 positive cells were counted manually over all the fields of view. The proportion of dying cells was obtained by calculating the ratio of this number relative to the total cell number. Motility was determined by tracking the nuclei of Nuclight positive cells. As above, the nuclei were segmented with the trained Cellpose model. ImageJ TrackMate plugin [42] was applied on the segmented images. For each stack, the velocities at every step were pooled together, regardless of the cell trajectory they belonged to. The median velocity was calculated from the pooled data set. Mean of the medians was then calculated for every condition.

Change in cell morphology was quantified by measuring the intensity of fluorescence of the cytoplasm of the GFP-tagged U87-MG. When a cell shrinks, the intensity level per pixel increases since it is the projection of the volume concentration of the GFP. To conduct this analysis, the GFP images were segmented using a Cellpose-trained neural network. The network was trained in a similar way to that used for nuclei detection. The raw GFP images were then multiplied by the binarised mask to obtain the distribution of intensities of every pixel belonging to cell bodies. This distribution was corrected by the background signal as follows: the mask was inverted, multiplied by the raw image. The median intensity of the resulting image was calculated. The median background intensity was subtracted from the cell intensity distribution. This correction was applied frame-by-frame. Negative values were filtered out.

### 2.9 Immunofluorescence and confocal imaging

Two hours post field stimulation, the cells were permeabilised for 15 min in a solution of PBS^+^*^/^*^+^ containing 4% PFA and 0.5% of Triton 100x. They were then fixed with a solution of 4% of PFA in PBS^+^*^/^*^+^ for 45 min. The permeabilisation step allowed to significantly decrease the fluorescence levels of the GFP in the cytoplasm of the cells and of the Nuclight stain. Vimentin was stained with a mouse monoclonal antibody (Vimentin V9 Ref. sc-6260, Santa Cruz Technology) at dilution 1:50 and a goat anti mouse CY2 secondary antibody (Jackson Immunology, ref. 115225146) at dilution 1:200. Actin was stained using Alexa Fluor Plus 647 Phalloidin (Invitrogen, ref. A30107) at dilution 1:400 following provider’s recommendations. Finally, the nuclei were stained with Hoechst 33342 at dilution 1:1000. These concentrations of antibodies allowed to get a signal that strongly dominates the remaining fluorescence of the cytoplasm and of the Nucligth stain.

The stained samples were imaged with confocal microscopy (LSM 880, Zeiss) at either 63x and 40x magnification, using oil immersion objectives. In-depth images of the samples were acquired with a spacing of 0.3 *µ*m. MDs were imaged at the same altitudes using the reflected light at 488 nm.

### 2.10 Traction force microscopy

Cells were seeded at 4000 cell/cm^2^ on the polyacrylamide hydrogels designed for traction force microscopy (TFM). Image acquisition was performed using a 60x oil immersion objective (NA 1.4) 2 h and 5 h after the cells have been exposed to the magnetic field, or, for the control condition, 26 h and 29 h post seeding. Stacks of images of the fluorescent beads were acquired in the vicinity of the surface of the hydrogel with 0.3 *µ*m spacing in order to compensate for any vertical drift of the surface. In parallel to capturing images of the fluorescent beads, phase contrast and GFP images were captured to visualise the cells and the MDs. The latter appear either black or bright green on the GFP images when in contact with cells, while they are more visible on phase contrast images, when they lay in the surrounding area. At the end of the experiment, the cells were detached using 0.5% trypsin-EDTA (Trypsin-EDTA 10 x, Gibco, Thermo Fisher Scientific, Gennevilliers, France, ref. 15400) for one hour. This allowed to image the relaxed state of the gel, or reference state.

The quantification of the hydrogel deformation field is detailed in a previous publication [43]. In brief, the deformation field is obtained by measuring the displacement of fluorescent beads located in the close vicinity of the surface. This is achieved by comparing the beads position while the cell is present and pulling on the gel, the strained state, to the cell-free hydrogel, the reference state. A reference image is selected from the reference stack, that is the closest to the surface of the gel. A home-made registration algorithm was used to compensate for the in-depth and lateral drifts of the strained image by comparing the beads position in regions of the strained state that are far from the cells (and therefore where a reduced deformation of the substrate is expected) to the reference image. Once the images were corrected for drifts, the deformation field was quantified using Kanade–Lucas–Tomasi pyramidal optical flow algorithm as described in [43]. Between 45000 and 52000 fluorescent beads were tracked on images whose size is of the order of 2000 × 2000 pixels. Four levels in the pyramid were used, thus working with subsampling the original image with new pixel sizes of respectively 64, 16, 4, 1 times the original pixel area. The size of the interrogation windows used to track the displacements were respectively 40^2^ , 30^2^ , 20^2^, 20^2^ pixels, the unit pixel size referring to that of the level of the pyramid. The calculation of force field was performed using Fourier transform traction cytometry [44]. The noise level of the traction forces was quantified from the force field out of the cells. To do so, masks were drawn based on the GFP and phase contrast images to differentiate cell body from background. Since the U87-MG have the tendency of overlap and form small groups, the forces were analysed per field of view and not per cell, as it was not possible to single out individual cells within colonies. Then a mask of areas devoid of MDs and cells was drawn for each image and the distribution of forces out of cell bodies and adhered particles was calculated. The 0.025 and 0.975 quantiles of the distribution of the *x* and *y* components of the traction stresses were calculated. The upper value of the quantile was used to filter the data measured in cells: only values that exceeded this threshold were used for further analysis. An example of this filtering process is reported in supplementary Figure S1.

### 2.11 Statistical analysis

Every assay was performed in triplicate, with cells coming from different passages in order to yield three independent experiments (n=3). Mean and standard deviations were calculated over these three values.

One sample Student’s t-tests compared to 1 were used whenever we calculate ratios or normalised values (cell detachment or proliferation), with 0.05 significance level to reject the null hypothesis. As we expect the magnetic stimulation to damage cells, we used one-tailed t-tests to test the null hypothesis when comparing the positivity of PI-staining in cells that have been stimulated by the magnetic field to the control, unstimulated cells. Similarly, one-tailed Student’s t-tests were used to assess the significance of the contribution of MDs vibration to the increase in the proportion of PI-positive cells compared with the 0 Hz condition. In all other cases, two-tailed Student’s t-tests were used. For all tests, a significance level of 0.05 was chosen to reject the null hypothesis.

For the analysis of morphological changes, the corrected distributions of pixels values was pooled between the three independent experiments, condition per condition. The cumulative distribution function (CDF) was then calculated over the whole set of data.

Regarding the motility measurement, the median velocity per condition was calculated from the distribution of velocities of every cells displacement in each independent experiment. Means of the medians were used to asses any statistical difference with the control using two-tailed t-tests, as previously described. The data of the three experiments were then pooled to calculate the general distribution per condition.

To evaluate any significant difference in cellular traction forces between conditions, the median values of the filtered distribution of the cellular traction forces was calculated per field of view. The mean of the medians was then calculated per experiment. Two-tailed t-tests were used to compare to the control condition, consisting of cells loaded with MDs but not exposed to the magnetic field. Moreover, the distribution of filtered force amplitudes were pooled by condition to calculate the associated CDF.

## 3 Results

### 3.1 The magnetic particles exert oscillating torques and compressive forces when actuated by a rotating Halbach array

Using FEM, we calculated the magnetic torque that arises when a magnetisation is submitted to the oscillating field (Fig. 4). This models a MD that is immobilized, and whose magnetisation is saturated and remains so under field oscillations. As the torque value depends on the relative orientation between the MD magnetisation and the local magnetic field, we have considered the three limiting cases where the saturated magnetisation is along the *x*, *y* or *z* direction. Focusing on the central region of the array where the cells are grown, the largest torques are experienced when the MD magnetisation is within the *xy* plane, with an amplitude of about 15 nN·*µ*m (Fig. 4b,d). Vertical-pointing magnetisation along *z*-axis results in a much reduced magnetic torque (Fig. 4f). Because the MDs are highly anisotropic in shape, the magnetic torque **T** only transfers into a mechanical torque when it is parallel to the flat plane of the particle, the *xy* plane. And for instance, *T_z_*torque induces magnetisation rotation within the *xy* plane, but no mechanical torque. Considering now the orbital motion of the magnets in the *xy* plane, the simulation shows that it induces either a variation in the magnetic torque amplitude when the particle is saturated along the *x* or the *z* axis, or a transfer of the torque from the *x* to the *z* axis while keeping its amplitude fairly constant when the particle is saturated along the *y* axis (Fig. 4). This simulation also indicates that the order of magnitude of the torque is in the nN·*µ*m range, therefore being sufficient to reorganize and alter cell cytoskeleton [45] or disrupt cell membrane [46], but not sufficient to induce a thermal effect [40].

**Figure 4:**
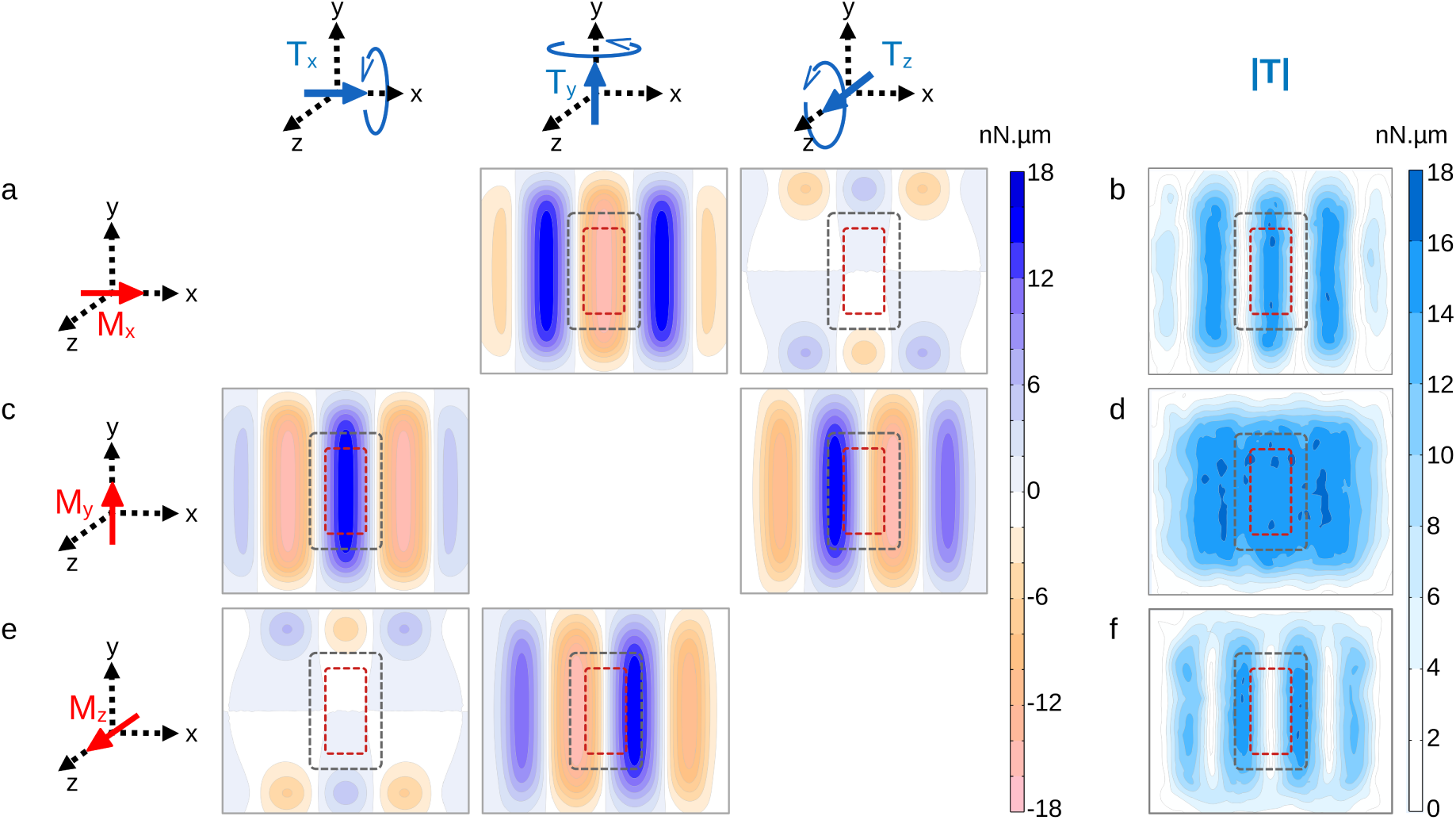
2D maps of the magnetic torques applied by the Halbach array to immobilised particles positioned 4 mm above the magnets and whose magnetisation is saturated. The maps are organized in a table. Different orientations of the magnetisation, illustrated by a red arrow on the schematics on the left of the table, are shown in rows. The blue arrows in the schematics on top of the table illustrate components of the rotational motion possibly induced by the magnetic torque **T**. The last column shows |**T**|, the amplitude of the torque vector. (a–b) The magnetisation is oriented along the *x* axis. (a) Components along *x*, *y* and *z* axis and (b) amplitude of the magnetic torque. The torque is predominantly aligned along the *y* axis. (c–d) Same with the magnetisation being oriented along the *y* axis. In this geometry, the torque components along the *x* and *z* axis have similar amplitudes. (e–f) Same with the magnetisation being oriented along the *z* axis. The torque is predominantly aligned along the *y* axis. The red and the grey dashed rectangles show the regions of interest, resp. in static regime and when the array is set in orbital motion.

In addition to the magnetic torques, the finite elements computation also shows that the Halbach array exerts forces on the MD. As shown in Fig. 5 the forces in the *xy* plane are below 1 pN. On the other hand the vertical, downward force is larger, of about 3.5 pN. Such force amplitude can be expected to trigger ion channel opening, but is not sufficient to rupture membranes [47]. However, the actual forces might exceed the values reported here in the case where particles aggregation occurs in cells. These clusters are expected to give rise to forces of larger magnitude, in proportion to the number of particles in the aggregate. Field-induced translational forces could then become sufficient to provoke cellular damages, such as impairment of the integrity of the cytoskeleton or disturbance of protein assemblies [48].

**Figure 5:**
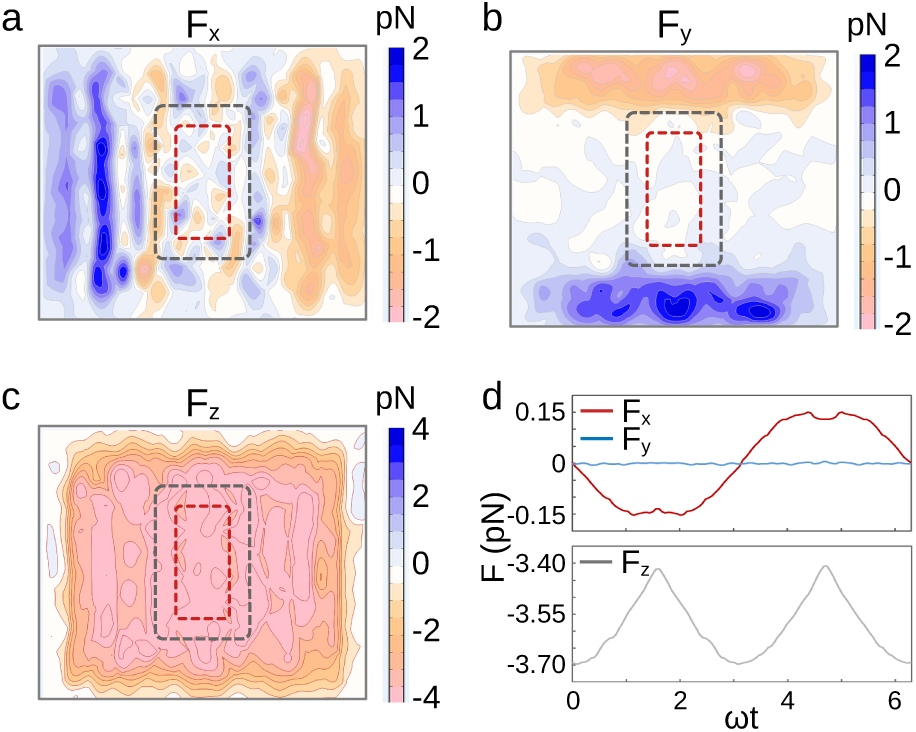
(a–c) 2D maps of the translational forces on particles with saturated magnetisation along the local field. The particles are positioned 4 mm above the magnets. The red and grey dashed rectangles show the regions of interest, resp. in static regime and when the array is set in orbital motion. (d) Time variation of the translational forces for a particle positioned at the center of the red dashed rectangle and saturated along the *x*-direction. *ω* is the pulsation of the orbital movement of the array of magnets.

### 3.2 At equal particles load, physiologically soft extracellular environment enhances the adverse effect of stimulation over cell detachment, viability and proliferation

U87-MG cells were first exposed to the magnetic microdiscs at a concentration such that the cells are loaded with an average of 500 particles per cell. Since cells grown on a stiff or on a soft substrate do not capture the same amount of microdiscs [32], the volume concentration of particles was adapted to each condition (see Materials and Methods). In a previous work, we had shown that this concentration of microdiscs does not significantly alter the viability of U87-MG cells on both substrates [32]. The cells were then exposed to the magnetic field, of amplitude 250 mT. The frequency of the rotation of the Halbach array was assayed between 0 Hz (static field) and 10 Hz.

Our first observation is that the magneto-mechanical stimulation has more adverse effects in cells grown on the soft substrate. For instance, a significant proportion of cells, of about 40%, has detached of the soft support 4 h after the 10 Hz, 20 min long magnetic stimulation (Fig. 6a) (p = 0.01). This is in contrast to glass (p = 0.01), where all the cells remain adhered. Similarly, the loss of viability is more pronounced in cells grown on 10 kPa substrate than on glass (Fig. 6b,c) (t-test, respectively 4 h and 24 h post stimulation: p = 0.02, 0.009 (Ctrl), p = 0.03, 0.08 (0 Hz), p = 0.04, 0.002 (2 Hz), p = 0.01, 0.03 (10 Hz)). Nevertheless, field application has an effect in both culture conditions. In particular, we note that exposure to the static field alone for 20 min (0 Hz) triggers an increase of the amount of cells entering death processes in the first hours following the stimulation (p = 0.01 (glass), p = 0.02 (10 kPa)). Superimposed alternating field leads to additional significant damages, but only in cells grown on the 10 kPa support (p = 0.03 at 10 Hz). This effect persists 24 h after the field has been released (Fig. 6c, p = 0.02). This is in contrast to glass on which setting in vibration microdiscs that exert a static pressure does not induce additional alteration in cells compared to a stimulation consisting of a static pressure only (p = 0.5 at 10 Hz). And finally, we observe that the magnetic stimulation halts cell proliferation, but again this happens only on the soft substrate (Fig. 6d) (t-test respectively on glass and 10 kPa: p = 0.001, 0.007 (Ctrl), p = 0.03, 0.23 (0Hz), p = 0.0008, 0.13 (2 Hz), p = 0.01, 0.11 (10 Hz)).

**Figure 6:**
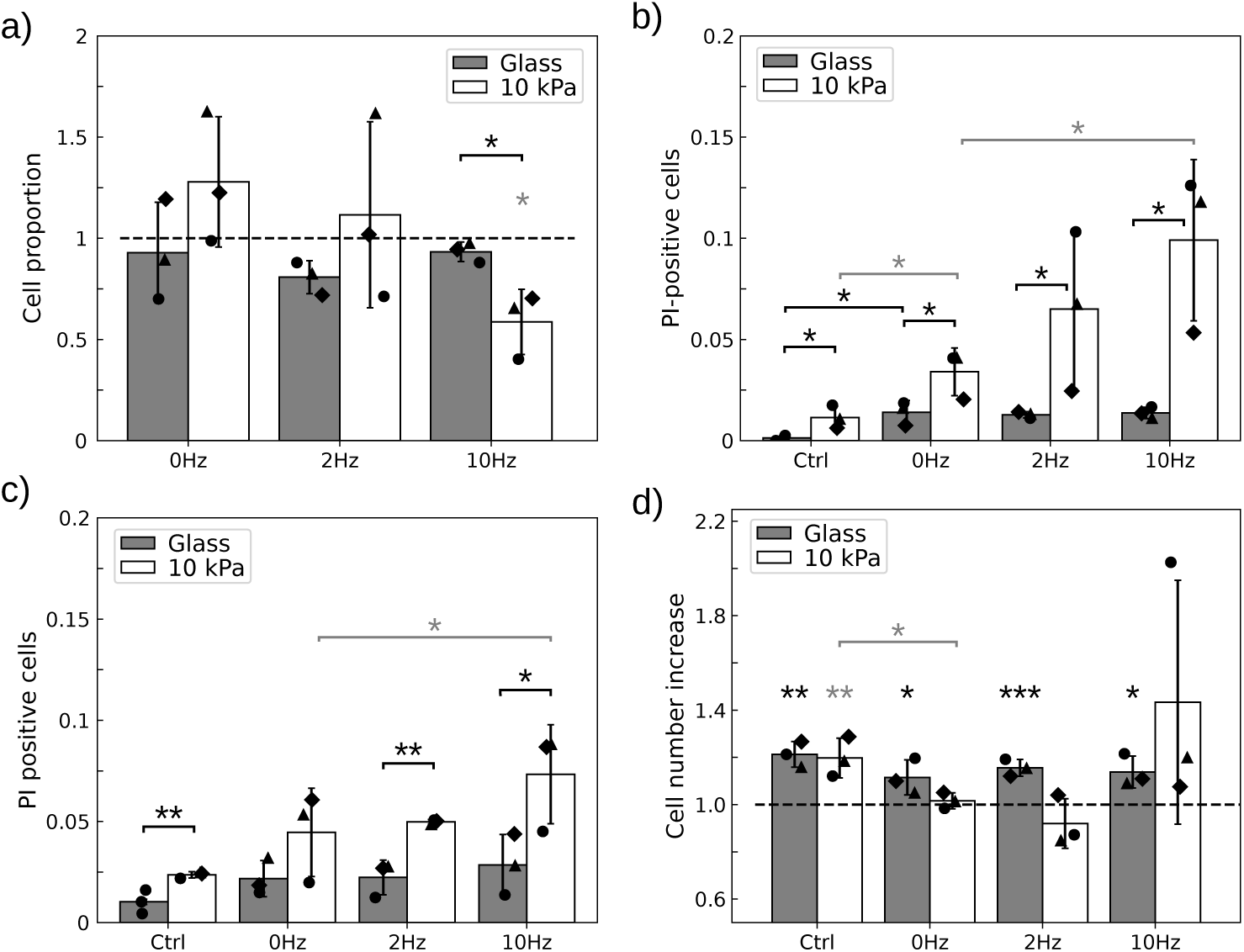
Field application perturbs cells more when grown on 10 kPa substrate. a) Proportion of remaining cells respective to control 4 h after field application at 3 different frequencies, for the two substrates: glass and 10 kPa polyacrylamide hydrogel both coated with 1 *µ*g/cm^2^. b – c) Application of the magnetic field increases the proportion of PI-positive cells (b) 4 h and (c) 24 h after field application. Vibrating particles have a significantly stronger detrimental effect than those submitted to a static field on the soft substrate. d) Cell proliferation is halted after field application in cells grown on 10 kPa substrate, in presence of either a static or an oscillating field. Black symbols show independent experiments (n=3). Error bars represent standard deviation of the means. *, **, *** denote significance with respectively p *<* 0.05, 0.01, 0.001. Gray stars refer to statistics for the soft substrate condition while black stars refer to the glass substrate. Only significant differences are labelled.

As a whole, we conclude that U87-MG cells loaded with an average of 500 MDs/cell are more affected by the magnetic stimulation when they are grown on a physiologically soft substrate than on a substrate with an out-of-physiology stiffness. The magnetic actuation of the MDs results in cell detachment, halt in cell proliferation and loss of viability. An important observation is that while the static field, and the resulting vertical pressure exerted by the particles, is enough to induce damages in the cells, vibrating particles trigger additional detrimental effects that are only visible in cells grown on soft substrates although the average load of particles per cell is identical in both culture conditions.

### 3.3 At lower particle load, cells grown on 10 kPa exhibit significant frequency-dependent halt in proliferation, and transient reduction in viability and motility

Our goal then is to better understand how the actuated particles may alter cell normal fate. Loading cells with an average of 500 MDs leads to cell detachment for cells grown on the soft substrate (Fig. 6a). This inevitably results in cell selection, with only the least particle-laden cells or the most robust phenotypes being selected. This selection then influences, and may bias, subsequent cell viability or proliferation experiments. And indeed, we observe that the dispersion of the data is much larger in the experiments conducted on the soft substrate than on glass where the cell number does not decrease after treatment. We attribute this difference to the selection in favour of the cells least affected by the mechanical disturbance. As detachment may depend on the state of the neighbouring cells, therefore on the local cell density, the state of the remaining cells is susceptible to be more variable. We confirmed this hypothesis by comparing the dispersion in the proportion of PI-positive cells through out the substrates in each independent experiment (Fig. S2).

When the load is reduced to 250 particles/cell, the stimulation does not induce significant cell detachment, thus avoiding to select specific phenotypes (Fig. 7a, p*>*0.3 for all conditions). Then we can evaluate the impact of the magnetic stimulation on the heterogeneous cell population, loaded with variable amount of particles as would occur in conditions where cells cannot be discarded from the experiment (like in vivo).

**Figure 7:**
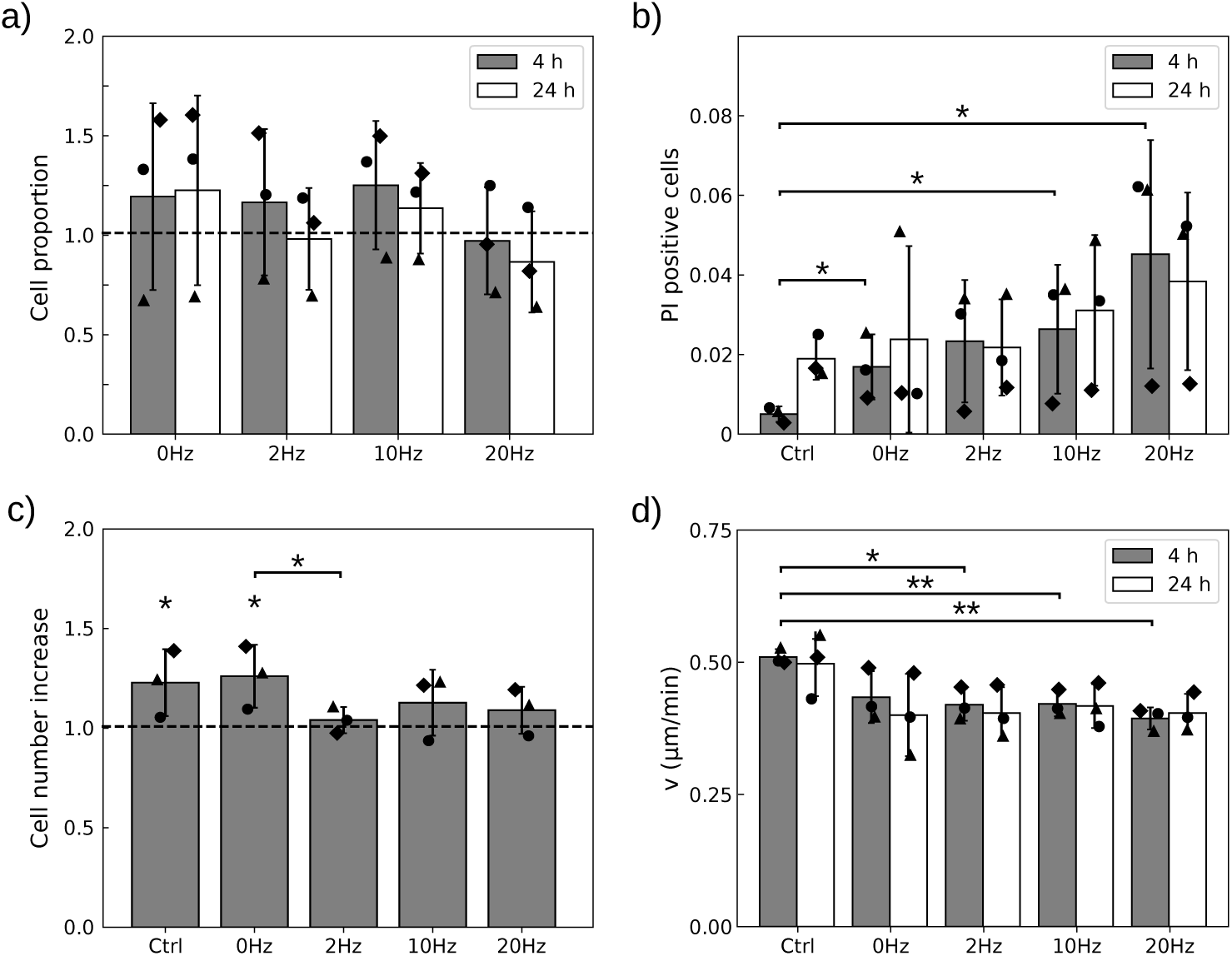
With a mean load of 250 MDs/cell, field application alters cell viability, proliferation and motility. a) Cells remain attached to the 10 kPa substrate when treated with a static pressure (0 Hz) or vibrating particles. The proportion is calculated relative to the control in the absence of field. b) The proportion of PI-positive cells increases 4 h but not 24 h after field application relative to the control. c) Cell proliferation is halted after field application in cells submitted to an oscillating field but not a static one. d) Cell migration is on averaged slowed down 4 h after the treatment but recovers 20 h later. Black symbols show independent experiments (n=3). Error bars represent standard deviations of mean (a – c) of median (d) values. * and ** denote significance with respectively p *<* 0.05 and 0.01. Only significant differences are labelled.

With this lower dose of MDs, we thus report on the response of the whole cell population, having heterogenous load of particles and therefore variable intensities of perturbation. Figure 7b shows that 4 h post treatment, cell viability is significantly altered at all frequencies, including the static treatment (0 Hz). Although the effect seems to increase with frequency, a frequency-dependence could not be assessed due to the large variability in cell responses (Anova test: p = 0.35). Nonetheless the frequency of the treatment has a significant impact on cell proliferation and motility (Fig. 7c,d). Proliferation as well as motility are impaired only when the particles vibrate: with this load of particles, static compression (0 Hz) is not sufficient to alter these functions, although 20 min of vibration at any of the assayed frequencies does. Nevertheless, the impact of the mechanical stresses exerted by the particles on cell viability and motility decreases in time. None of these read-outs remained significant 24 h post treatment. Thus we conclude that the most obvious persistent effect of the stimulation is to stop cell proliferation for at least 24 h.

### 3.4 Cell morphology on soft substrate is persistently altered at both microdiscs loads assayed but only when vibrated

As shown above, mechanical perturbations of the cells by the actuated MDs have consequences whose persistence in time depends on the cellular function considered. The ones we focused on, viability, proliferation or migration are key concerning cancer evolution but they result from the integration of many highly regulated biochemical reactions. In contrast, cell shape is a lower level read-out in terms of biological complexity, that can be predictive of cell outcome [49]. We therefore focused on cell morphology and analysed how the mechanical perturbation would impact it. By carefully looking at cell shape dynamics, we observed that in some conditions, after an immediate collapse, some cells were spreading again few hours later (Fig. 8a). We took benefit from the fluorescence of the GFP tagged U87-MG cells to quantify the change in morphology, collapsed cells having a larger fluorescent signal per pixel (Fig. 8b). We therefore measured the fluorescent signal of every pixel associated to a cell and plotted the cumulative distribution function (CDF) of the normalized, background-corrected, intensity. An example of such a cumulative function is shown for cells grown on the soft substrate, bearing an average load of 500 MDs per cell and observed 4 h after magnetic stimulation (Fig. 8c). It shows that after stimulation, the intensity of fluorescence is shifted toward larger values compared to the control condition, meaning that cells have adopted a more rounded morphology. In all conditions, we observed that the CDF departs from the control condition toward larger fluorescent intensity (Fig. S3). We quantified this observation by calculating the Wasserstein distance (*d_w_*) between the control condition (particle-laden cells unexposed to the magnetic field), and those submitted to a static or alternating field (Fig. 8d). The Wasserstein distance provides here a metrics of the integrated distance between the two CDFs. Comparing the Wasserstein distances calculated 4 h and 24 h post treatment allowed to assess whether the cells recover a morphology closer to the control or evolve toward an even more rounded shape, a priori associated to dying processes (Fig. 8e). As a whole this analysis highlights that the shape of cells grown on the soft substrate is persistently altered when they are exposed to a 10 Hz or 20 Hz stimulation, at the two doses of MDs assayed. This is in contrast to glass where the Wasserstein distance decreases in time at 10 Hz, suggesting that the cells recover progressively their morphology.

**Figure 8:**
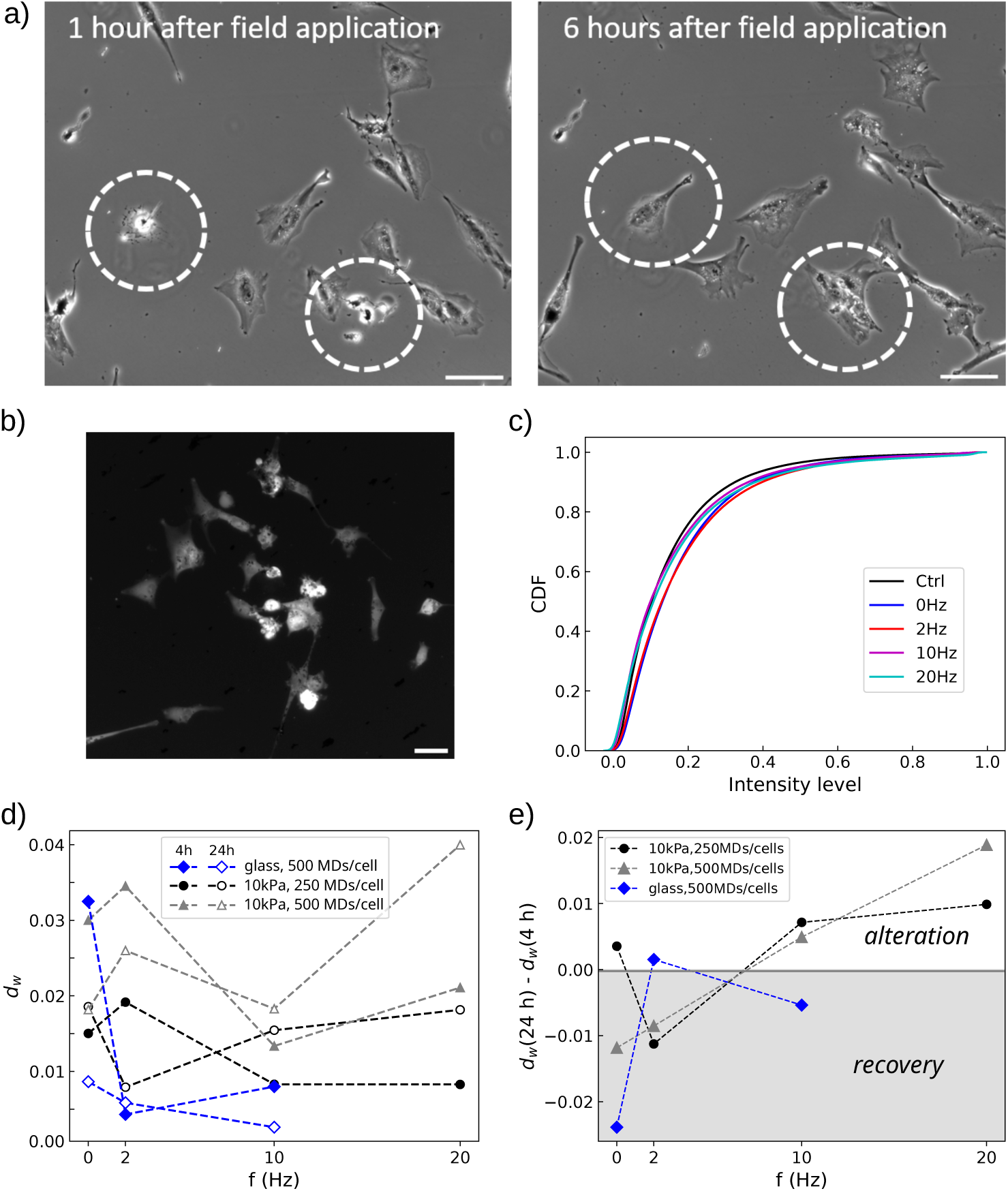
a) Cells undergo morphological changes due to the field-induced particle actuation, collapsing to a round shape. The morphological change is reversible when the cell is not dying, as highlighted by the cells inside the dashed circles. Scale bar 100 *µ*m. b) The shrinkage of the cells causes an increase of the fluorescence of the pixels in the cell coming from the reduced volume. Scale bar 50 *µ*m. c) Cumulative distribution function (CDF) of the fluorescence intensity per pixel of the cells 4 h post treatment, when the cells are grown on 10 kPa substrate and are loaded with 500 MDs/cell. d) Wasserstein distance (*d_w_*) between control conditions and cells submitted to the magnetic field measured 4 h or 24 h post treatment, for different culture conditions (glass and 10 kPa substrate) and density of MDs (250 or 500 MDs/cell). e) The variation of the Wasserstein distance in time informs on the evolution of cell morphology compared to control.

Figure 8d also shows that the static field has a short term impact on cell morphology in all culture conditions, that is more pronounced at larger concentration of MDs, in consistence with the fact that the static field generates a downward mechanical pressure whose amplitude is proportional to the number of particles per unit surface. Nevertheless, we observe that the morphology of cells grown on glass evolve toward their unperturbed morphology faster than those grown on the soft substrate at equal particle load (Fig. 8e). The evolution of the Wasserstein distance with frequency seems to be non monotonic in all conditions, and differs between cells grown on glass or on the soft substrate (Fig. 8d). For instance addition of a mechanical vibration at 2 Hz seems to have a large initial impact on cell morphology in cells grown on the soft substrate, but cells progressively recover a spread morphology few hours later. This is in contrast to the effects of vibrations at 10 Hz or 20 Hz, that appear more persistent.

### 3.5 The microdiscs interact with the actin and vimentin cytoskeletons

We observed that cell morphology is differentially altered as a function of the stiffness of the substrate and the frequency of the mechanical perturbation. Cell shape closely relates to the organisation of the cytoskeleton, which differs whether cells are grown on a soft or a stiff substrate [50, 51] (Fig. S4). We therefore questioned whether the mechanical vibrations of the particles, that enhance the detrimental effects of particle actuation on cell viability, proliferation or migration compared to a compressional stress alone, could lead to visible damage of the cytoskeleton organization.

In previous studies [11, 32], we and others had shown that in the absence of surface functionalization, the MDs are internalized in cells. TEM imaging [11] as well as DIL staining [32] did not highlight any membrane around the particles, suggesting that they could enter cell body through a translocation process [52]. And indeed, many of them can be seen in the cytoplasm of the GFP-tagged U87-MG cells (Fig. S5). We thus imaged the actin and the vimentin cytoskeletons of cells loaded with an average of 250 MDs that had been submitted to a compressional stress alone or a combination of compressional and vibrational stresses. Both cytoskeletons were immuno-stained 24 h post treatment and compared to particle-laden cells that were not exposed to the magnetic field (Fig. 9).

**Figure 9:**
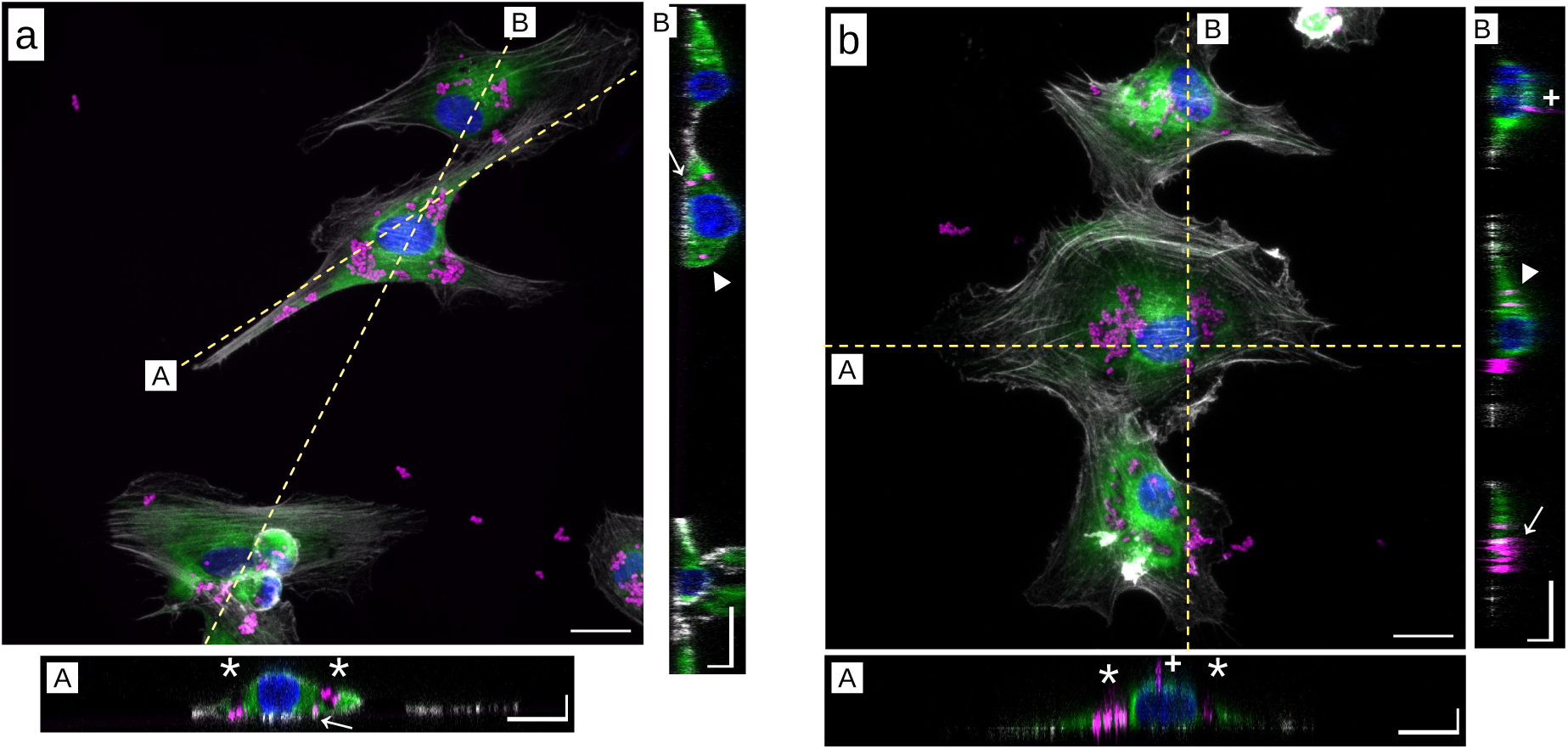
Representative snapshots of immunostained U87-MG cells exposed to the microdiscs for 24 h in the absence of magnetic field. The cells are grown on (a) glass or on (b) 10 kPa substrate. Actin is shown in grey, vimentin in green and nuclei in blue. Bottom and right hand side panels show in-depth cross sections of cells acquired along the labelled dashed lines. The white stars point toward microdiscs trapped in void regions of the vimentin cytoskeleton. The white triangles (resp. arrows) point toward particles embedded into the vimentin (resp. actin) cytoskeleton. Plus (+) symbols show microdiscs bound to the membrane. In-plane bar: 20 *µ*m. In-depth bar: 5 *µ*m.

Confocal imaging provided some information on the localization of the MDs relative to the cytoskeleton. We observed that the particles are positioned either at the cell membrane, into the vimentin network or in contact with actin fibres, or that they are trapped into vacuoles devoided of vimentin and actin that are presumably related to autophagy (Fig. 9). Of note, U87-MG cells are cancer cells with highly active autophagy processes even in the absence of particles [53]. And we confirmed that these vacuoles embedded into the vimentin cytoskeleton are also found in the absence of particles, although they are more frequent in their presence (Fig. S4). Few depletion of actin filaments were observed in cells exposed to a 10 Hz stimulation, while none were visible in the vimentin cytoskeleton (Fig. S6). No such actin depletion was visible in the control condition. Nevertheless, these events were rare and hardly visible. They could not be quantified to conclude about a differential alteration of actin cytoskeleton when the frequency is varied.

### 3.6 The vibrations of the microdiscs alter cell contractility

The optical analysis of the actin and vimentin cytoskeletons provides us information on the location of the particles relative to them but this analysis lacks resolution to conclude about the impact of the vibrational movement of the particles on their integrity. For this reason, we quantified the mechanical stresses the cells exert on the substrate. These stresses inform on cell contractility [54], a read-out of the integrity of the cytoskeleton.

We conducted this analysis by both looking at the cells-averaged contractility and the subcellular force patterns. We compared the contractility of cells loaded with an average of 250 particles/cell and exposed to static (0 Hz) or alternating fields (2, 10 , 20 Hz) to two control conditions: MD-free cells and MDs-laden cells, both in the absence of magnetic field (Fig. 10). The control without particles was performed to identify a potential perturbation of cellular forces coming from the internalised particles, in the absence of magnetic field.

**Figure 10:**
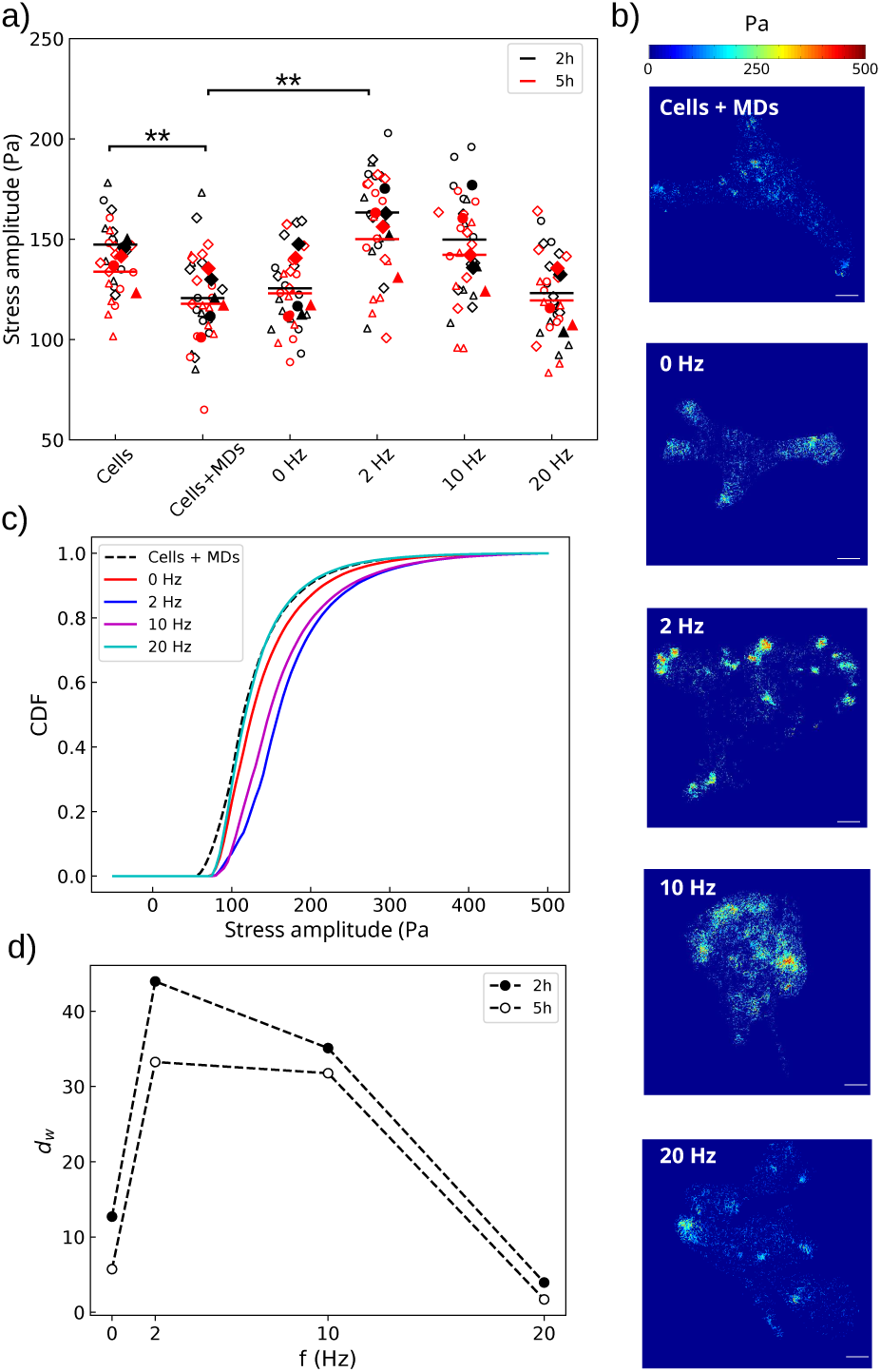
The mechanical stimulation enhances cell contractility. a) Amplitude of the cellular stresses transmitted to the substrate. Empty symbols show median values for each field of view, while filled symbols are the mean of the medians per experiment. Different shapes indicate independent experiments (n=3). The horizontal bars are the means of the replicates. Significance was calculated against the control condition Cells + MDs with two-tailed t-student test. Black (resp. red) colour shows results and statistics 2 h (resp. 5 h) after stimulation. ** denotes significance with p *<* 0.01. Only significant differences are labelled. b) Representative traction force patterns, 2 h post treatment. The control condition consists of particle-laden cells not exposed to the magnetic field. Scale bar 25 *µ*m. c) Cumulative distribution function (CDF) of the stress amplitude for the different mechanical perturbations. d) The Wasserstein distance (*d_w_*) shows that the 2 Hz and 10 Hz treatments have the greatest impact on cell contractile capabilities, but that this impact tends to decrease in time.

Amplitude maps of the mechanical forces per unit surface the cells transmit to the substrate were calculated from the displacements of nano-scaled fluorescent polystyrene beads embedded into the hydrogel substrates as described in Materials and Methods. The stress map was restricted to the inside of the cellular contour. The outer signal, at the exception of that coming from non-internalised magnetic MDs lying on the substrate, was used to determine the noise level of the measurement. The inner signal could thus be filtered and only stresses above the noise level were considered (Fig. S1). Data were acquired 2 h and 5 h after magnetic stimulation to analyse the immediate evolution of cellular forces following the mechanical perturbation.

We first observed that, in the absence of magnetic stimulation, cells having no particles inside exert significantly larger stresses than those loaded with 250 particles/cell (Fig. 10a, p = 0.008). This evidences that although the MDs were shown to be inert concerning cell metabolic activity, viability, proliferation or migration at this dose of particles [32], they do perturb a very basic regulator of many cellular functions, the contractile machinery. Secondly, while we do not observe any significant variation of the cellular stresses after the particles have exerted a static compression (p_0_*_Hz_* = 0.71), we observe that the 2 Hz treatment results in a significant increase of the amplitude of the stresses (p_2_*_Hz_* = 0.008). Increase of the frequency of the vibration however leads to less significant outcome, the effect being fully lost at 20 Hz (p_10_*_Hz_* = 0.11, p_20_*_Hz_* = 0.83). This suggests that the vibrations of the MDs only alter cell contractility in a narrow range of frequency, of few Hz. Looking at the median values of the traction forces, the perturbation appears transient, the significance of the effect being lost 5 h post treatment (p_2_*_Hz_* = 0.08 ,p_10_*_Hz_* = 0.16, p_20_*_Hz_* = 0.90). However, analysing the median values alone might lead to incomplete results, seen the difference in the distribution of the force points through the cells after they were submitted to an alternating field or not (Fig. 10b). Large amplitude traction points were often observed in cells submitted to the stimulation, in contrast to the control. This qualitative observation was made more quantitative by comparing the cumulative distribution function of the force amplitudes (Fig. 10c). Consistently, we observed a positive shift of the CDFs at 2 and 10 Hz relative to the non stimulated cells loaded with MDs, meaning that these two conditions generate local forces of larger amplitude. In contrast, the 20 Hz stimulation did not induce any change in the amplitudes of the local cellular stresses compared to those of the control condition. Calculation of the Wasserstein distance relative to the non-stimulated control condition additionally showed that the alteration in cell contractility persists 5 h post treatment but only when the cells are stimulated with 2 Hz or 10 Hz (Fig. 10d).

## 4 Discussion

Previous studies using conventional 2D culture plates have demonstrated that low-frequency, field-induced vibration of micrometric disc-shaped particles can elicit cell death at remarkably high rates, ranging from 60% to 90% [13, 15, 17, 55]. Notably, Kim et al. [13] and Leulmi et al. [55] functionalized these discs to target the cell membrane, achieving comparable results across various cancer cell lines. The initiation of apoptotic pathways was attributed to an elevated influx of calcium ions, driven by the magnetic-induced stimulation [13]. Additionally, studies have reported instances where internalized particles, when actuated, trigger cell death [15, 17, 56], suggesting the potential of this approach for innovative anti-cancer therapies. However, translating these promising in vitro findings to more complex environments, such as 3D artificial scaffolds or in vivo models, has proven challenging [15, 25, 26]. Among the factors differentiating conventional culture substrates from 3D substrates and in vivo tissues, the mechanical properties of the extracellular environment are particularly significant. Notably, culturing cells on 2D soft substrates has been shown to better preserve cell phenotypes and enable transcriptomic expression profiles that closely resemble those observed in vivo [36].

Motivated by the discrepancies in cell outcomes observed between conventional plastic plates and more biomimetic environments, this study examines the role of extracellular matrix softness in shaping cell fate following the actuation of magnetic microdiscs. Specifically, we seek to determine whether the mechanical properties of the extracellular environment modulate the cellular response to the microdiscs-induced mechanical perturbations. Our experimental setup involves superimposing compressive forces and shear stresses coming from particles actuation. The soft extracellular environment is mimicked using a 50 *µ*m-thick layer of polyacrylamide hydrogel coated with fibronectin, a protein of the extracellular matrix. Glass substrates coated with the same amount of fibronectin serve as controls for conventional 2D culture. This protocol enables to discriminate the impact of the extracellular matrix softness from other cues, such as chemical interactions of the cells with the surface.

Here we employed U87-MG cell line, a well-established in vitro model of gliobastoma cells. The response of these cells to intracellular particle actuation has been analysed in several former studies, using conventional culture plates [13, 15, 17, 56]. They all concluded to the efficiency of the vibrating particles to induce cell death. When performing similar analysis with cells grown on a soft substrate, of 10 kPa, our main observation is that the cells are even more sensitive to the magneto-mechanical stimulation compared to those grown on stiff substrates. This suggests that even when used at a non toxic dose, the particles may trigger significant responses in cells grown on substrates with physiologically relevant mechanical properties at lower doses than previously anticipated in plastic plates.

More specifically, our first key finding is that the mechanical perturbation exerted by the static pressure of the MDs is sufficient to induce cell death and limit proliferation, with this effect being more pronounced in cells grown on a soft substrate (Fig. 6). Our second key finding is that superimposing vibration onto the compressive force significantly enhances treatment efficacy in cells grown on the soft substrate, unlike those grown on glass (Fig. 6). Although it was not possible to assay vibration alone in this setup, the lack of significant difference in cell death rates between static compression and combined compressive and vibrating stimulations at a lower dose of MDs (Fig. 7) suggests that both stimulations play complementary roles in compromising cell integrity.

A plausible mechanism underlying this phenomenon is that the compressive forces exerted by the particles in the cell body impair the integrity of cellular membranes and the cytoskeleton. This is supported by our observation that the MDs interact with both cell compartments [32] (Fig. 9, Fig. S6). As demonstrated by Kim et al., membrane bending may activate mechanosensitive ion channels, initiating cell death programs [13]. Additionally, compressive stresses in the order of tens of kPa (nN/*µ*m^2^) can rupture the cell membrane [46], a range achievable when several hundred MDs aggregate. Similar compressive forces are also in principle sufficient to alter the integrity of membrane-bound organelles such as autophagosomes and lysosomes, leading to the release of their acidic content [57].

The differential response of cells grown on a soft or a stiff substrate may be attributed to the softer rheological properties of cells grown on soft substrates [58–60]. This is associated to a reduced content of actin stress fibres [61] (Fig. S4). The adaptation of actin cytoskeleton organisation to the stiffness of the extracellular matrix is well-documented [62–64] and correlates with changes in cellular rheological properties [65]. In our study, as the MDs are internalized by the cells (Fig. 9), their motion is constrained by the mechanical properties of the cell interior, with stiffer compartments impeding particle vibration [40]. This may explain why adding vibration to the compressive force has no additional effect on cells grown on glass, whereas on the 10 kPa substrate, vibration significantly enhances cell death and detachment (Fig. 6). Another factor contributing to the resilience of cells grown on hard substrates to mechanical perturbation is the different organization of vimentin intermediate filaments and their interaction with actin and microtubule cytoskeletons. The vimentin cytoskeleton exhibits visco-elastic properties [66]. It thus dissipates mechanical energy [67] and provides mechanical support to various cellular functions, such as trafficking and adhesion [68]. Moreover, vimentin can directly cross-link to actin and microtubules and stabilise them [63]. And finally, vimentin is upregulated in cells grown on stiff substrates [61], which may provide an additional protection to the cells against intracellular mechanical perturbations. Consistently, we qualitatively observe separated vimentin and actin cytoskeletons in cells grown on the 10 kPa substrate, while they are intertwined in cells grown on glass (Fig. S4).

Soft substrates, by allowing the translation of MD forces and torques into noticeable deformation of intracellular compartments, thus appear as an interesting platform to study the influence of local mechanical vibrations in cells grown in mechanically relevant environments. Working with a reduced load of 250 MDs/cell on average allowed to limit cell detachment and phenotype selection. At this load, we observe that the static pressure exerted by the particles is sufficient to transiently alter cell viability and morphology, but not proliferation or motility (Fig. 7, 8). In contrast, superimposing a vibration onto this compressive force significantly impairs cell proliferation and motility, while it does not significantly enhance the loss of viability compared to a static pressure alone. The impact on cell morphology is also more persistent when the particles are set in vibration (Fig. 8). Although we could not clearly attribute the impact of vibrations to a disorganisation of the actin cytoskeleton, we found that particles vibrations indeed alter cellular traction forces, an indirect readout of cytoskeleton integrity (Fig. 10). We observed that this alteration is frequency dependent, being maximal at 2 Hz and 10 Hz while it is lost at 20 Hz. This observation is consistent with our theoretical analysis predicting a cut-off frequency above which the vibration of the MD is dampened [40]. At low frequency, the alteration of contractility levels and the concentration of force points around focal points (Fig. 10) hints at the laceration of actin filaments. Similar findings were reported by Kumar et al. [69] who showed that cutting single actin filaments using a laser ablation technique leads to a transient increase in cell contractility and localisation of force point around focal adhesions. Indeed, we expect the MD vibration to apply torques in the nN.*µ*m range (Fig. 4). The excursion angle of the oscillating magnetic field being close to *θ*_0_ ≃ 1rad, the oscillatory motion of the MD explores a fraction of the arc *Rθ*_0_ ≃ 0.6µm, depending on the visco-elastic properties of the intracellular compartment [40]. Therefore the force exerted by one vibrating MD on the actin network is likely larger than a few nN, resulting in a local pressure in the order of a few kPa. This value is well above experimental results measuring in vitro actin networks breakage for stresses of 0.1 Pa [45]. Thus MD vibration is capable of disrupting the actin cytoskeleton, unlike vimentin network which resists much larger strain and stress before breaking [45]. And indeed, we consistently could not observe any depletion of vimentin in the vicinity of vibrating MDs, unlike actin (Fig. S6).

At 2 Hz and to a lesser extend at 10 Hz, the alteration in cell contractility persisted for 5 h post-treatment. Within this frequency range, we also observed persistent alteration in cell morphology and proliferation 24 h post treatment (Fig. 7 and Fig. 8). These findings support the idea that vibrating MDs alter cell contractility, a primary marker of cell fate, consistent with numerous studies correlating cell contractility levels with proliferation or migration capabilities [70–72]. However, this correlation did not hold when comparing contractility and cell morphology evolution in cells exposed to 20 Hz orbital field (Fig. 8d). While the vibration of the MDs had no noticeable effect on cell contractile capabilities, cell shape was significantly altered, evolving in time toward more rounded shapes. These cells also exhibited the highest death rate (Fig. 7b). While the limited impact on cell contractility can be understood by the reduced deformation of the intracellular material at this larger vibration frequency [40], yet it results in more severe damages to cells. This suggests that the loss of viability at higher frequency is likely due to the way the particles disorganise the actin (or vimentin) cytoskeleton and other cellular compartments, rather than the amplitude of the perturbing oscillation and stress.

## 5 Conclusion

Overall, our study demonstrates that the mechanical properties of the extracellular environment significantly influence the efficacy of magneto-mechanical stimulation by micrometric magnetic discs. Glioblastoma cells grown on soft polyacrylamide substrates mimicking the stiffness of fibrotic tissues were more sensitive to treatment than those grown on stiff substrates at equal particle loads. This enhanced sensitivity was attributed to stiffness-induced cytoskeletal reorganisation and the resulting changes in cellular rheology. Furthermore, we showed that superimposing vibration onto compressive forces enhances treatment efficacy in a frequency-dependent manner. At low frequencies (2–10 Hz), the mechanical stimulation markedly affected the actomyosin machinery, indicating that the cytoskeleton is a primary target. At higher frequencies (*>*10 Hz), alterations in morphology, proliferation, and motility did not correlate with actin contractility impairment, suggesting the involvement of additional mechanisms. These findings underscore the potential of magneto-mechanical stimulation as both a standalone and adjunct nanomedicine strategy for glioblastoma treatment.

## 6 Authors’ contributions

AV performed the experiments under the supervision of RM and AN. AV and SS implemented pipelines to analyse the data. RM performed the FEM modelling and simulations. AV, RM and AN analysed and curated the data. AV, RM, BD and AN designed the experiments. BD, RM and AN acquired the project funding, conceptualised the study and developed the methodology. AV, RM and AN wrote the initial draft which was reviewed and edited by all authors.

## Acknowledgements

This work is supported by the French National Research Agency in the framework of the “Investissements d’avenir” program (ANR-15-IDEX-02) and by the french Renatech network. This project received help from MuLife imaging facility, which is funded by GRAL, a program from the Chemistry Biology Health Graduate School of University Grenoble Alpes (ANR-17-EURE-0003). The authors acknowledge the support of the French Renatech network.

