## Supplementary figures for "Reducing Glioblastoma Cell Aggressiveness via Static and Dynamic Magneto-Mechanical Stimulation with Vortex Microdiscs on Substrates of Physiological Stiffness"

[1]Univ. Grenoble Alpes, CNRS, CEA/LETI-Minatec, Grenoble INP, LTM, Grenoble F-38000, France.
[2]Univ. Grenoble Alpes, CEA, CNRS, Spintec, Grenoble F-38000, France.

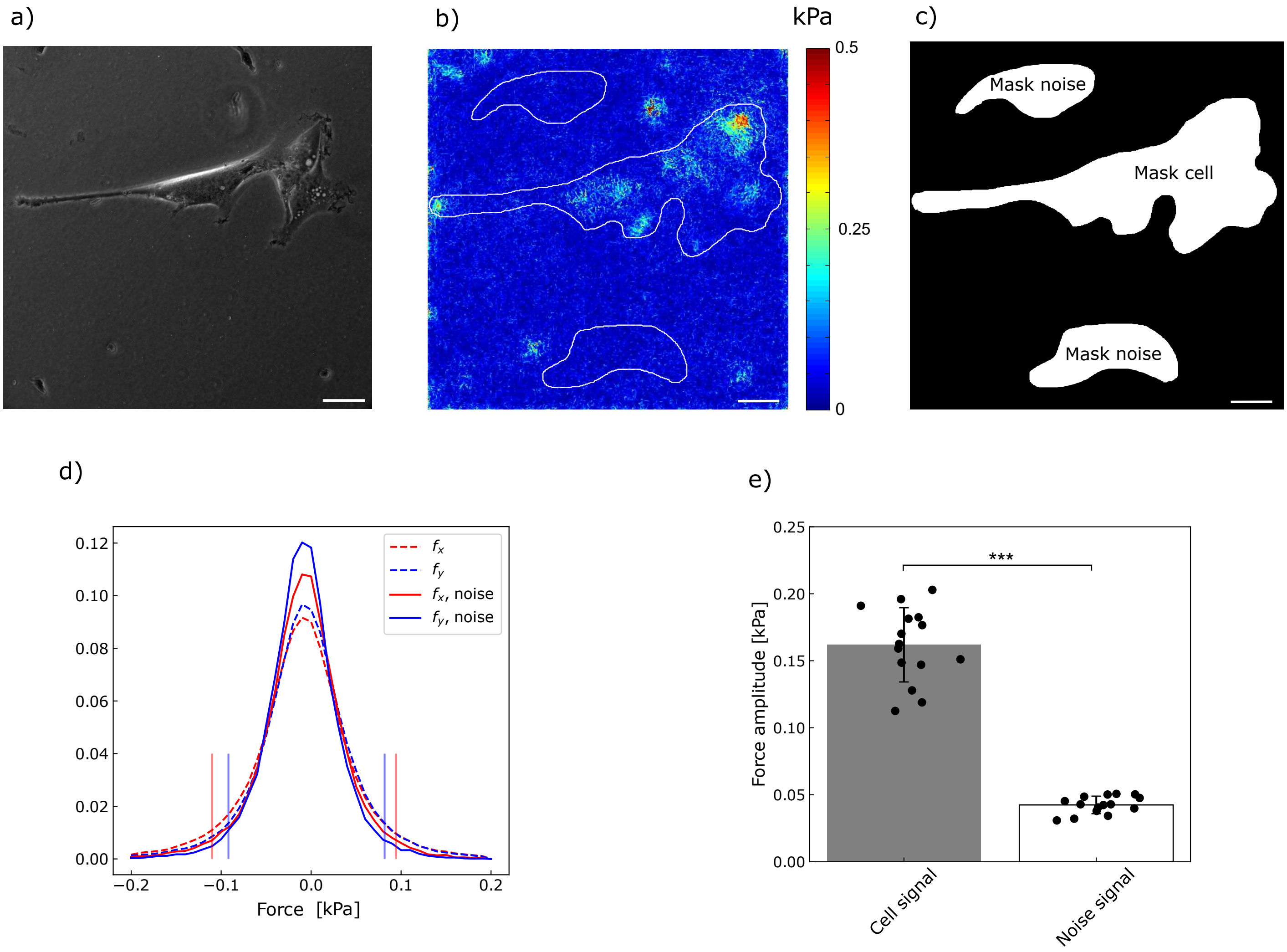


**Figure S1** (a - c) Example of the filtering process used to calculate the noise level on the amplitude of cellular forces. a) Phase contrast image of the field of view with U87-MG cells. b) Force amplitude map and the contours of the masks used to extract the forces inside the cell contour and the noise level. Scale bar 25 $\mu$m. c) Binary masks used to extract the force field exerted by the cells (labelled "Mask cells") and to quantify the noise level (labelled "Mask noise"). Aggregates of MDs laying on the surface experience adhesion forces and generate substrate deformation, therefore they are discarded from the quantification of the noise level. d) Distribution of the in-plane traction force components, $f_x$ and $f_y$, of the cells (dashed lines) and the ones of the noise (solid lines). Vertical lines indicate the quantile 0.025 and 0.975 of the noise level for $f_x$ (red) and $f_y$ (blue). d) Mean of the medians of traction force amplitudes of the cells after filtering the noise level, in one representative experiment. Bars indicate the standard deviation. Significance was calculated by T-student test (n=15). *** denotes p-value $<$ 0.001.

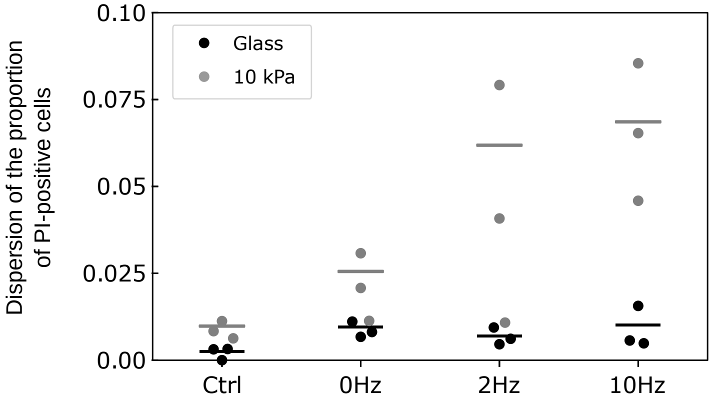


**Figure S2** The proportion of PI-positive cells shows a larger variability across the sample when the cells are grown on 10 kPa substrate compared to glass support. The dots show the dispersion throughout 5 fields of view of 1.3 x 1.3 mm$^2$ chosen randomly (n=3). Bars show the total dispersion per condition, calculated as $\sigma^2 = \frac{1}{nn_i-1}\left(\sum_{i=1}^{3}(n_i-1)\sigma_i^2 + \sum_{i=1}^{3} n_i(\bar{w_i}^2 - \bar{w}^2)\right)$ with $\bar{w_i}$ and $\sigma_i$ being respectively the average and the dispersion of the proportion of PI-positive cells in each experiment, $\bar{w}$ the average of the experiments, $n$ the number of experiments and $n_i$ the number of fields of view in the $i^{th}$ experiment (n=3, $n_i = 5$ for all $i$).

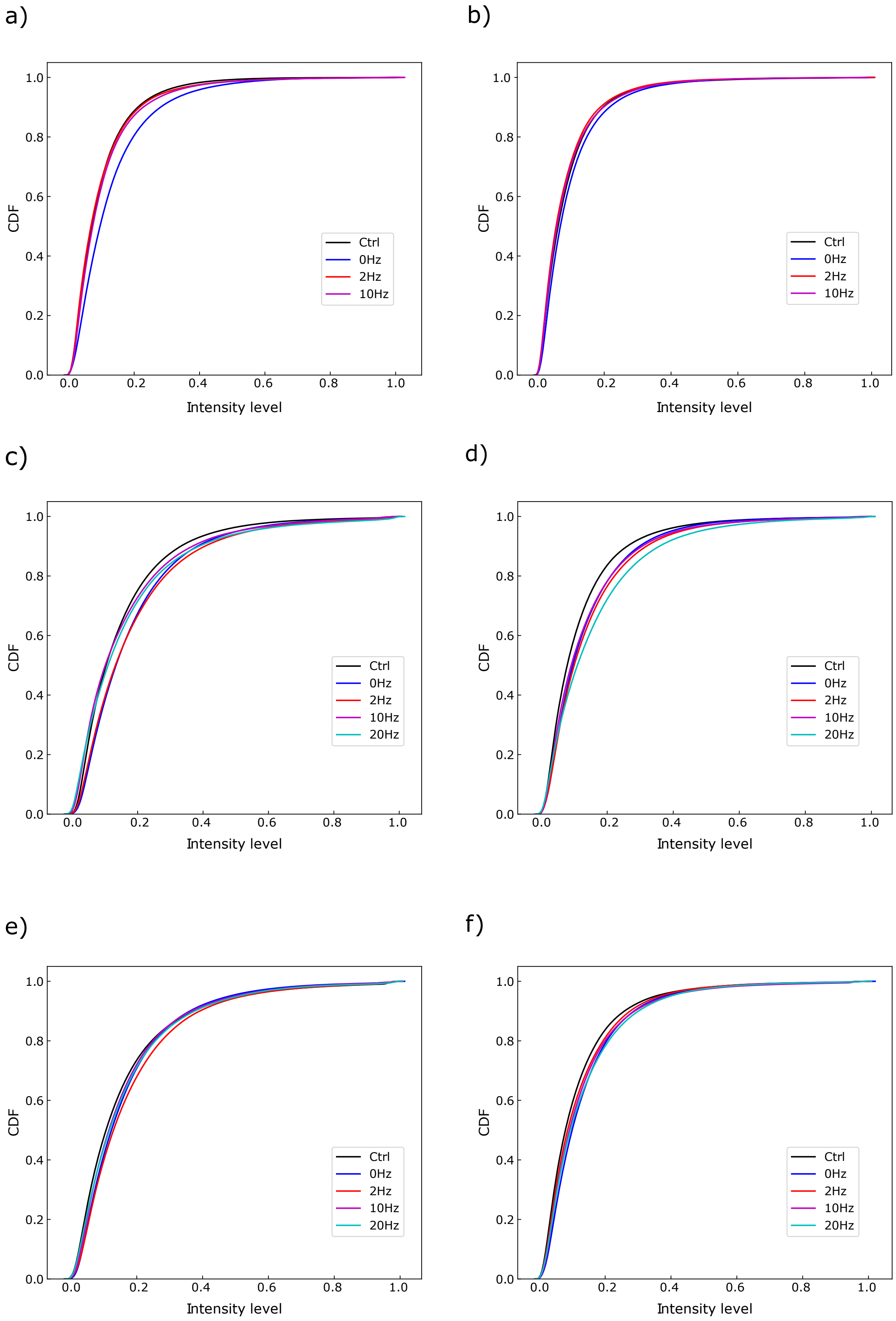


**Figure S3** Cumulative distribution function (CDF) of the fluorescence intensity per pixel of the cells. (a-b) cells grown on glass loaded with 500 MDs/cell, 4 and 24 h post treatment respectively. (c-d) cells grown on 10 kPa loaded with 500 MDs/cell, 4 and 24 h post treatment respectively. (e-f) cells grown on 10 kPa loaded with 250 MDs/cell, 4 and 24 h post treatment respectively. The fluorescence intensity per pixel is normalised to the maximum intensity of the image.

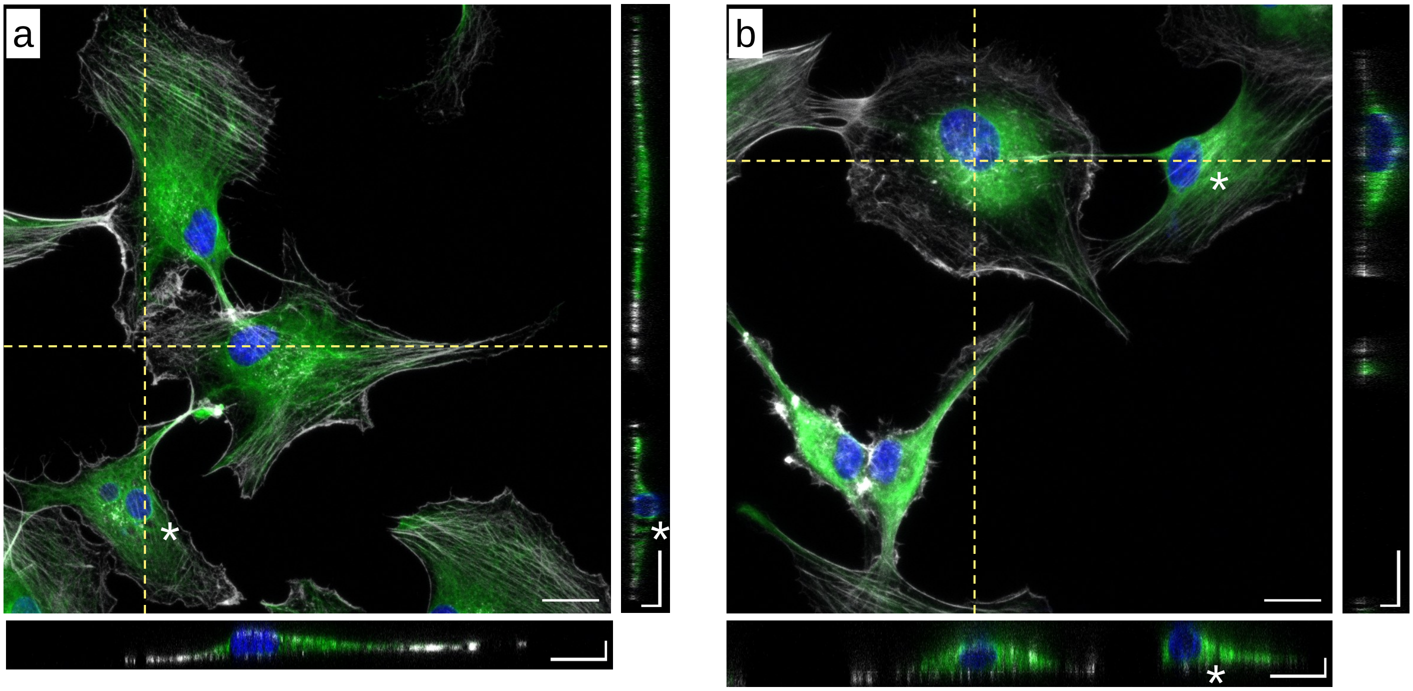


**Figure S4** Representative snapshots of immunostained U87-MG cells grown on (a) glass or on (b) 10 kPa substrate. Actin is shown in grey, vimentin in green and nuclei in blue. Bottom and right hand side panels show in-depth cross sections of cells acquired respectively along the dashed lines. The white stars point toward holes in the vimentin cytoskeleton. In-plane bar: 20 $\mu$m. In-depth bar: 5 $\mu$m.

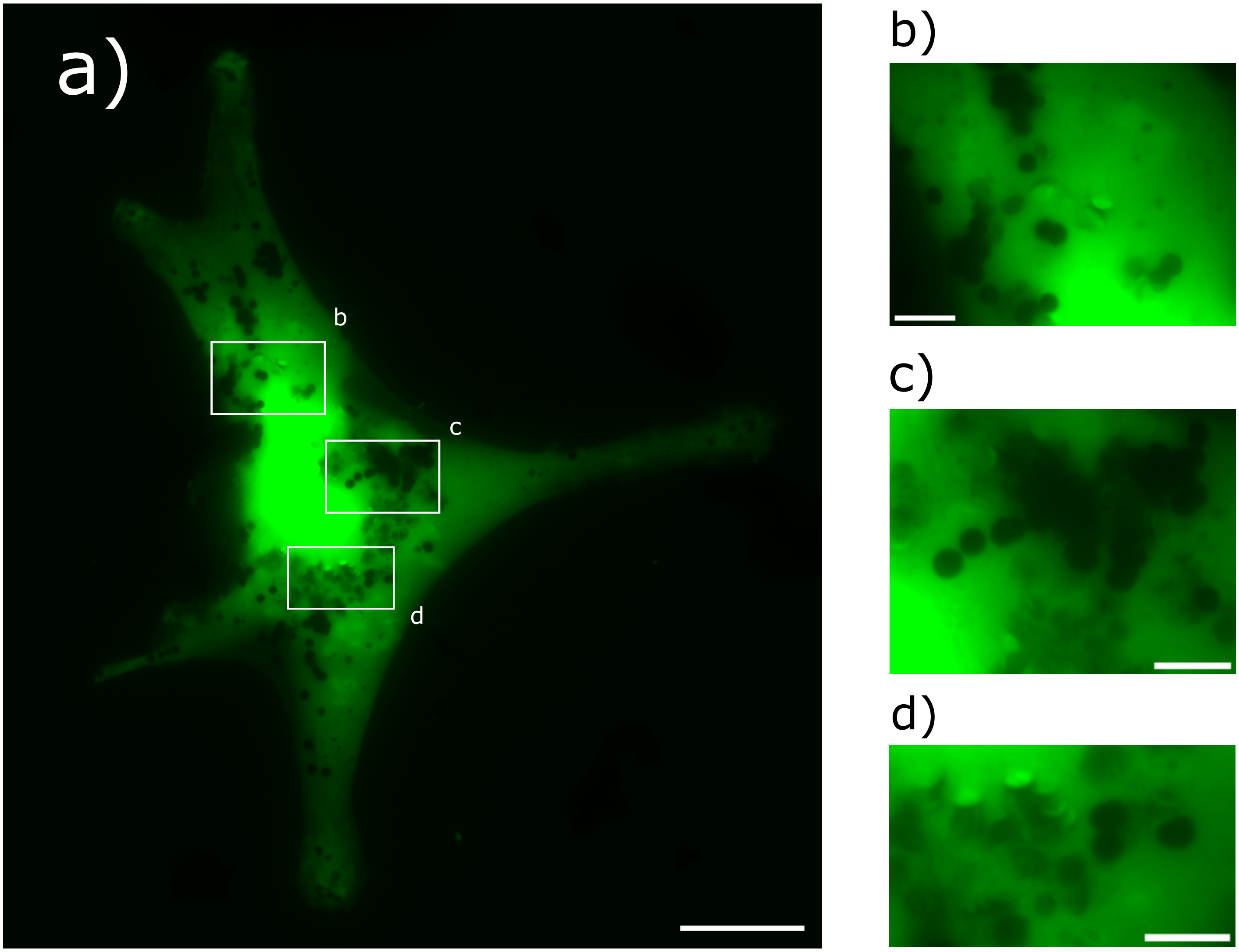


**Figure S5** The MDs are internalized by U87-MG cells and have various orientations in the absence of magnetic field. (a) The particles appear dark when they are oriented parallel to the culture plate, when they screen the fluorescence of the cytoplasm. They appear bright when they are tilted relative to the plane of the culture plate as they reflect the fluorescence of the cytoplasm. Bar: 25 $\mu$m. (b-d) Zoom of the regions enclosed in the white square in (a). Bar: 5 $\mu$m.

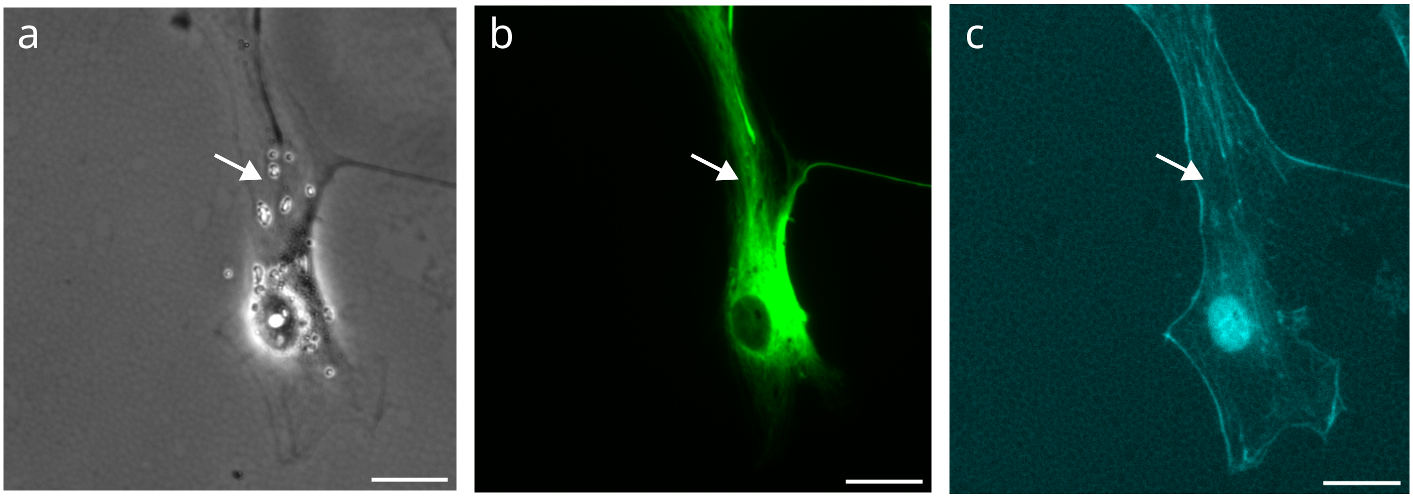


**Figure S6** The vibrations of the particles remodel the actin cytoskeleton. U87-MG cells grown on a 10 kPa substrate were loaded with 250 MDs/cell and exposed to 10 Hz stimulation. Images are captured 24 h post treatment. (a) Phase contrast image. (b) Vimentin cytoskeleton. (c) Actin cytoskeleton. Arrows indicate areas around MDs de-voided of actin cytoskeleton, while vimentin cytoskeleton is unaltered. Scale bar: 25 $\mu$m.
